# Prolonging Genetic Circuit Stability through Adaptive Evolution of Overlapping Genes

**DOI:** 10.1101/2023.02.27.530340

**Authors:** Jennifer L. Chlebek, Sean P. Leonard, Christina Kang-Yun, Mimi C. Yung, Dante P. Ricci, Yongqin Jiao, Dan M. Park

**Affiliations:** Biosciences and Biotechnology Division, Lawrence Livermore National Laboratory, Livermore, CA 94550

## Abstract

The development of synthetic biological circuits that maintain functionality over application relevant timescales remains a significant challenge. Here, we employed synthetic overlapping sequences in which one gene is encoded or “entangled” entirely within an alternative reading frame of another gene. In this design, the toxin-encoding *relE* was entangled within *ilvA*, which encodes threonine deaminase, an enzyme essential for isoleucine biosynthesis. A functional entanglement construct was obtained upon modification of the ribosome binding site of the internal *relE* gene. Using this optimized design, we found that the selection pressure to maintain functional IlvA stabilized the production of burdensome RelE for over 130 generations, which compares favorably with the most stable kill-switch circuits developed to date. This stabilizing effect was achieved through a complete alteration of the mutational landscape such that mutations inactivating the entangled genes were disfavored. Instead, the majority of lineages accumulated mutations within the regulatory region of *ilvA*. By reducing baseline *relE* expression, these more ‘benign’ mutations lowered circuit burden, which suppressed the accumulation of *relE* inactivating mutations, thereby prolonging kill-switch function. Overall, this work demonstrates the utility of sequence entanglement paired with an adaptive laboratory evolution campaign to increase the evolutionary stability of burdensome synthetic circuits.

## INTRODUCTION

For the past several decades, synthetic biologists have sought to genetically engineer microorganisms for a wide range of applications including therapeutics discovery and delivery, drug manufacturing, agricultural yields, biofuel production, mineral extraction, and waste degradation (1-8). For example, the microbial consortium that colonizes the rhizosphere of plant roots can be genetically engineered to enhance nutrient acquisition and drought resistance of agriculturally important crops (9,10). Such biotechnology applications require robust and stable expression of genetic circuits. Problematically, genetic circuit instability frequently originates from a fitness cost to the host due to leaky product toxicity (i.e., kill-switches) (11), host burden from adverse interactions with host components (12), or misallocation of resources (13-16). As a consequence, inactivating mutations accumulate and the resulting cells with disabled circuits rapidly outcompete the parent strain due to the relieved toxicity (17-20). For example, kill-switch circuits—which aim to control cell proliferation through regulated activity of a toxin—can fail when mutations arise that ablate the toxin’s function, allowing for lineages harboring non-toxic circuits to quickly overtake the population (21). Therefore, the development of tools that mitigate genetic instability while maintaining circuit function is necessary and central to synthetic biology.

Prior efforts to improve DNA sequence fidelity have focused on reducing background mutation rate (22,23), eliminating mutation-prone sequences (24), removing insertion sequence (IS) elements or avoiding them altogether through host selection (25,26), reducing burden of an engineered function (14), or increasing mutation surveillance and correction (27). However, the broad implementation of these genetic engineering efforts remains a challenge as it is difficult to predict which modifications are needed to achieve circuit stability in a given system *a priori*. Alternatively, laboratory evolution experiments have been used to optimize circuits by allowing for the unbiased selection of increased stability and performance (13,28). Previous work has sought to engineer systems that allow adaptive evolution (29) to maintain and stabilize the function of burdensome circuits, such as linking the expression of a toxic gene to an essential function (25,30,31) or dividing maintenance and production of a toxic gene between different members in a consortium (32). However, many of these stabilizing systems are complex, rely on multiple levels of redundancy, and have only been implemented in *Escherichia coli (33,34)*. Thus, more generalizable and effective methods are needed for improving genetic circuit stability, especially for sequence regions that are prone to mutational inactivation.

Recently, approaches using gene overlaps have been developed to enhance sequence stability (35-37). Gene overlaps occur naturally in many biological systems, especially those with high mutation rates and compact genomes such as viruses (38-42). Gene overlaps impose constraints on sequences and their evolution given that mutations can impact the function of both genes involved (39,43-45). A pioneering method to accomplish synthetic gene overlap is gene entanglement in which two genes are synthetically encoded (“entangled”) within the same DNA sequence but translated from different open reading frames (35). By entangling a gene-of-interest (GOI) with an essential gene, the evolution of the GOI can be constrained because mutations in the GOI may also be deleterious to the essential gene encoded in another frame. In one example in *E. coli*, entanglement of an amino acid biosynthetic gene *ilvA* with an essential gene *acpP* severely restricted the range of point mutations permissible within the overlap region of *ilvA* (35). While promising, the ability of synthetic gene overlaps to maintain genetic stability of engineered circuits remains untested.

Here, we demonstrate the use of sequence entanglement for improving genetic circuit stability, particularly for burdensome components that are prone to mutational inactivation. We assessed the feasibility of a proof-of-concept entanglement pair composed of a gene with high fitness cost (the toxin-encoding *relE*) and an essential gene (*ilvA*) to improve genetic stability of a toxin-based kill-switch circuit in the environmentally relevant microorganism *Pseudomonas protegens* Pf-5. Our findings provide insight into how gene entanglement alters the mutational landscape and evolutionary trajectory of synthetic circuits and showcase the ability of sequence entanglement to enable the use of natural selection to isolate cells with reduced fitness burden and more stable kill-switch function.

## METHODS AND MATERIALS

### Strains and culture conditions

*Pseudomonas protegens* Pf-5 was routinely grown in LB supplemented with kanamycin (Kan; 20 μg/mL), gentamicin (Gent; 15 μg/mL), or tetracycline (Tet; 25 μg/mL) when appropriate. Stellar competent *Escherichia coli* cells (Takara Bio) were routinely grown in LB supplemented with kanamycin (Kan; 50 μg/mL), gentamicin (Gent; 15 μg/mL), and carbenicillin (Carb; 100 μg/mL) when appropriate. All strains were grown overnight either shaking at ∼220rpm or statically at either 30°C (for *P. protegens*) or 37°C (for *E. coli*), unless otherwise noted. M9 minimal medium (Sigma) was supplemented with 1 mM MgSO_4_, 100 mM CaCl_2_, and 20 mM glucose.

### Construction of mutant strains

Primers (**Table S1**) were ordered from Integrated DNA Technologies (IDT) and constructs were PCR-amplified with Q5 High Fidelity polymerase (NEB). For all vectors, cloning was completed using InFusion (Takara Bio), according to the manufacturer’s instructions. RhaRS/P_*rhaBAD*_ and CymR/P_*cymR*_ (46) were added to the base vector (either pJUMP24-1A (47) modified with T24 terminator downstream of the cloning site or the miniTn7PuC18 vector (48), respectively) prior to cloning. All plasmids were sequence verified by Elim Biopharm or SNPsaurus. pJUMP24-T24-P_*rhaBAD*_ vectors were cloned into Stellar Competent *E. coli* using heat shock according to the manufacturer’s instructions and plasmids were maintained in LB with 50 μg/mL kanamycin.

Vectors were transformed into *P. protegens* via electroporation as described previously (48) and maintained in LB with 20 μg/mL kanamycin unless otherwise noted. In the *P. protegens* parent strain used in this study, CymR/P_*cymR*_-*relB* (the antitoxin) was integrated within the chromosomal attTn7 site (48) such that *cymR* expression was driven by the *lacIq* promoter, resulting in constitutive expression of CymR (**Fig. 1A**). All overnight cultures were prepared from glycerol stocks and grown in the presence of 0.5 mM cumate to relieve CymR repression and induce the antitoxin *relB*, except where indicated. See **Table S2 f**or a detailed list of all mutant strains used throughout this study.

**Fig 1.**
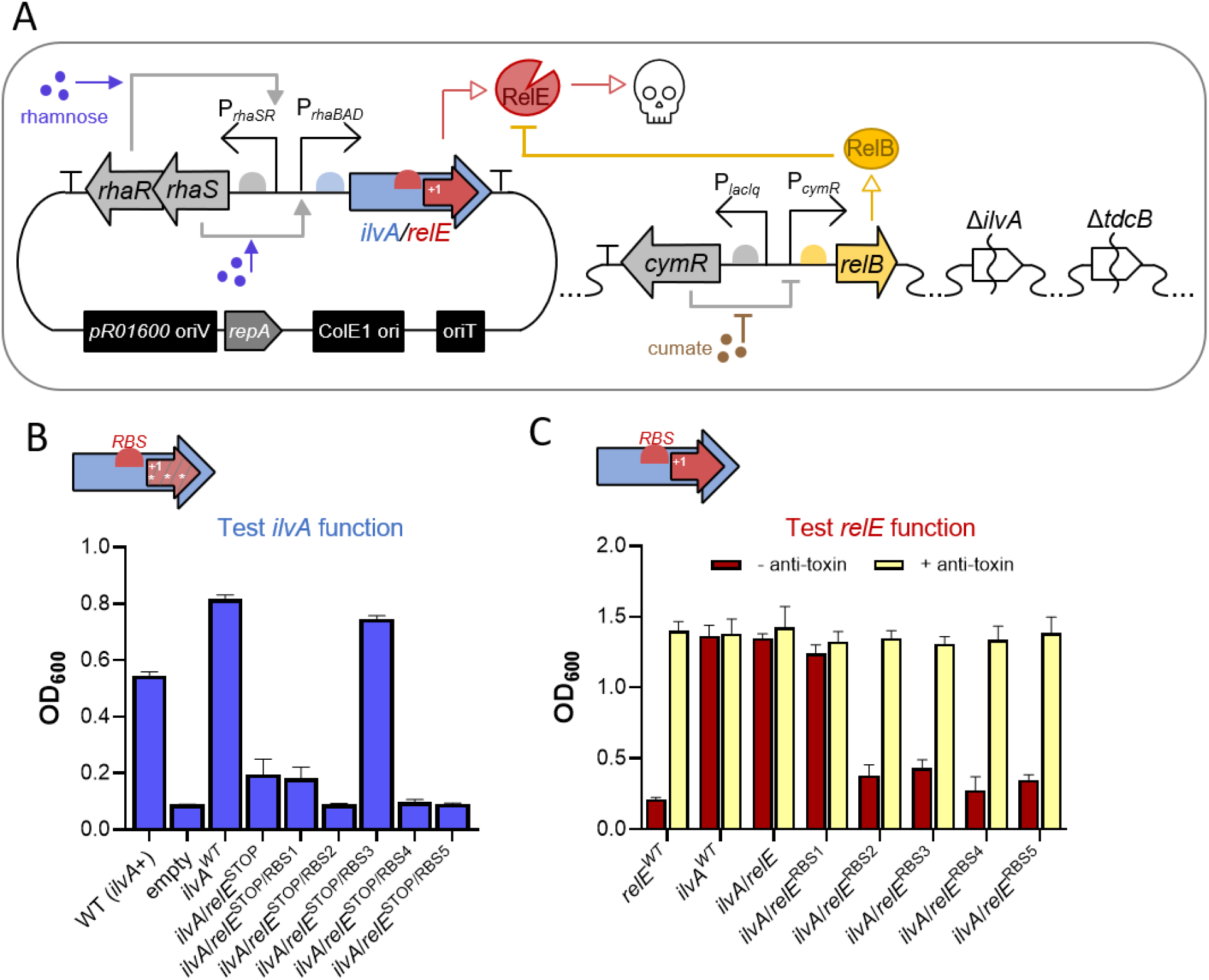
Internal RBS modifications improve functionality of *ilvA/relE*. **(A)** Diagram of genetic circuit components used in this study. *P. protegens Pf-5* was made auxotrophic for isoleucine via Δ*ilvA* and Δ*tdcB* chromosomal deletions (right). Addition of cumate induces antitoxin expression by relieving CymR repression of chromosomal P_*cymR*_-*relB* (right), while addition of rhamnose increases expression of *ilvA*/*relE* (or related alleles) through activation of plasmid-borne RhaR and RhaS, which activate P_*rhaBAD*_ (left). **(B)** To probe *ilvA* function, strains harboring *ilvA*/*relE*^*STOP*^ alleles containing different strength RBSs were grown in minimal medium without isoleucine and without addition of rhamnose. Stop codons in *relE* are represented by asterisks in the schematic. **(C)** Strains harboring *ilvA*/*relE* alleles with different strength internal RBSs were grown in rich medium to assess *relE* activity. To rescue growth, the antitoxin was induced by addition of cumate. For panels **B** and **C**, strains harboring *ilvA*/*relE* vectors are listed in order of increasing internal RBS strength (see **Fig. S5** for more details on RBS modifications). Growth is reported as OD_600_ after 15 h. Data are shown as the mean ± SD of 3 independent replicates.

In frame deletions of *ilvA* (PFL_5905) and *tdcB* (PFL_3098) in *P. protegens* were obtained by a two-step *sacB* counterselection procedure (49). Approximately 700 bp regions flanking the 5′ and 3′ regions of *ilvA* or *tdcB* were amplified using the primer sets described in **Table S1** and cloned into the HindIII and BamHI-digested suicide plasmid pNPTS138 using InFusion cloning. The pNPTS138-based deletion plasmids were transformed into *E. coli* MFDpir and conjugated into *P. protegens* Pf-5. Primary integrants were selected on LB containing 25 μg/mL kanamycin.

Counterselection for the second chromosomal crossover event, resulting in gene deletion, was selected for by overnight growth in LB media followed by plating on LB agar containing 3% sucrose. Deletions were confirmed by colony PCR and the Δ*ilvA* Δ*tdcB* strain was whole genome sequenced.

### Assessment of growth in liquid culture

Cells were grown overnight in LB +kanamycin +0.5 mM cumate and then washed twice in minimal medium. Cells were then diluted 1:25 in either LB or M9 minimal medium with kanamycin in Nunc™ Edge™ 96-Well flat bottom microplates (Thermofisher). The following concentrations of inducers were added when indicated: 0.001% (w/v) rhamnose to induce the expression of P_*rhaBAD*_ (50), 0.5 mM cumate to induce the expression of P_*cymR*_-*relB*, and 1 mM isoleucine. Assays were performed at 30°C with linear shaking and growth was kinetically monitored by measuring OD_600_ on an Agilent HTX plate reader.

### Western blot analysis

To assess threonine deaminase *(3xFLAG-ilvA)* production, strains were grown overnight in LB containing kanamycin. Strains were subcultured the next day 1:100 in fresh LB medium supplemented with kanamycin and 0.001% (w/v) rhamnose and grown shaking for 6 hours at 30°C. For each sample, cells were centrifuged and resuspended to an OD_600_ = 25.0 in B-PER™ Complete Bacterial Protein Extraction Reagent (Thermofisher) with 1mM PMSF and 38 μg/mL lysozyme and incubated at room temperature for 15 minutes. Samples were then mixed 1:1 with 2X SDS sample buffer [220 mM Tris pH 6.8, 25% glycerol, 1.8% SDS, 0.02% Bromophenol Blue, 5% *β*-mercaptoethanol] and then boiled for 10 minutes at 100°C. Then, 10 μL of each sample was separated on an Any kD™ Mini-PROTEAN® TGX™ Precast Protein Gels (BioRad). Proteins were electrophoretically transferred to a PVDF membrane using a Trans-Blot® Turbo™ Transfer System (BioRad) and then incubated with the primary antibodies α-FLAG polyclonal mouse (1:12,000) and α-RpoA monoclonal mouse (1:1,000) (Biolegend). Then, blots were washed and incubated with a α-rabbit and α-mouse secondary antibodies conjugated to horseradish peroxidase (HRP) and developed using Pierce ECL Western blotting substrate. Blots were imaged using a BioRad ChemiDoc instrument.

### Single time-point escape ratio

Single colonies of *P. protegens* Pf-5 were picked from LB-agar supplemented with kanamycin and 0.5 mM cumate and were inoculated in 3 mL of the LB medium supplemented with kanamycin and 0.5 mM cumate and grown with shaking at 30°C overnight. Cells were washed twice with minimal medium and serially diluted onto M9 minimal-agar containing kanamycin, 0.001% rhamnose, and 1mM isoleucine with or without 0.5 mM cumate (i.e., permissive or non-permissive conditions, respectively). Plates were incubated statically at 30 °C for ∼48 hours and then single colonies were picked and patched onto minimal medium-agar plates supplemented with kanamycin and 0.001% rhamnose either with or without 1 mM isoleucine. To calculate escape ratio, the colony forming units (CFU) per mL present on non-permissive medium was divided by the CFU/mL on permissive medium. For the condition without isoleucine, the ratio of colonies that survived on non-permissive plates without isoleucine was multiplied by the escape ratio from plates with isoleucine. To identify mutations causing escape, one colony per replicate was isolated from the patched plates for each growth phenotype; either they could only grow in the presence of isoleucine or under both conditions. The chromosomal P_*cymR*_*-relB* region of each colony was PCR amplified and the linear products were sequenced at Elim Biopharm or SNPsaurus. The plasmids of these colonies were purified via miniprep (Qiagen) and sequenced by SNPsaurus. A detailed list of mutations can be found in **Table S3**.

### Long-term evolutionary stability experiments

Single colonies of *P. protegens* Pf-5 strains were picked from LB-agar supplemented with kanamycin and 0.5 mM cumate and were inoculated into 5 mL of the minimal medium supplemented with kanamycin, 0.5 mM cumate, and 0.001% (w/v) rhamnose with or without 1mM isoleucine and grown shaking at 30°C for ∼6.6 generations (∼24 hours). Here, rhamnose was added to ensure sufficient *ilvA*/*relE*^RBS3^ expression for isoleucine production and cumate was added to induce sufficient RelB expression to minimize the toxic effects of RelE expression. Cultures were passaged with a 1:100 dilution into fresh medium at 24-hour intervals. To measure escape ratio, cells were washed once with minimal medium and serially diluted onto M9 minimal-agar containing the same inducers as grown in liquid culture, except with or without 0.5 mM cumate (i.e., permissive or non-permissive conditions, respectively). Plates were incubated statically at 30°C for ∼48 hours and then CFU/mL was counted. The escape ratio was calculated as the ratio of colonies on plates without cumate relative to total viable colonies on plates with cumate. Escape ratio measurements were taken at each 24-hour interval for 10 days (∼66 generations) or at a 5-day interval for 20 days (∼132 generations). The number of generations per day (n) was calculated as n = (log(N_t_) - log(N_o_))/log(2), where N_t_ is the final OD_600_, and N_o_ is the initial OD_600_. To identify mutations causing escape, colonies were isolated from the final passage of each separate lineage selected on permissive plates with 0.5 mM cumate. The chromosomal P_*cymR*_-*relB* region of each colony was amplified and the linear products were sanger sequenced at SNPsaurus. The plasmids of these colonies were purified via miniprep (Qiagen) and sequenced by SNPsaurus. A detailed list of mutations can be found in **Table S3**.

### GFP assay

Single colonies were grown in minimal medium containing kanamycin and 0.005% (w/v) rhamnose overnight at 30°C. The next day, cells were washed and resuspended to an OD_600_ = 1.0 in minimal medium. Fluorescence and OD_600_ was determined on an Agilent HTX plate reader with excitation at 500 nm and emission at 540 nm. Relative fluorescent units (RFU) are reported normalized to the OD_600_.

### Competition assay

Overnight cultures were grown in LB containing kanamycin and 0.5mM cumate and were washed twice and resuspended to a final OD_600_ of 1.0 in M9 minimal medium. Each culture was mixed in a 1:1 ratio with the parent strain. For each mixture, ∼10^5^ cells total were added to 5 mL M9 minimal medium supplemented with 0.001% rhamnose and 0.5 mM cumate and grown shaking at 30°C for 48 h. CFU/mL was determined at initial inoculation and after 48 hours by dilution plating on LB-agar containing kanamycin supplemented with either gentamycin or tetracycline. Competing strains were discerned by growth on either Gent or Tet containing plates, as the parent strain was Gent resistant/Tet sensitive, and the other strains were Gent sensitive/Tet resistant. Competitive indices were calculated as the CFU ratio of the evolved strain (CB strains 2, 3 and 4) over the parent strain after growth for 48 h divided by the CFU ratio in the initial inoculum *(51)*.

### Statistics

Statistical differences were assessed by Student’s t-test or one-way ANOVA tests followed by either a Dunnett’s or Tukey’s multiple comparisons post-test as indicated using GraphPad Prism software v. 9.5.0. For all escape ratio experiments statistical analyses were performed on log-transformed data. All statistical comparisons can be found in **Table S4**.

## RESULTS

### Post-Entanglement Modifications Enhances Functionality of Entanglement Pair

In order to assess gene overlaps as a method for improving the sequence fidelity of synthetic gene circuits, we selected a previously developed but moderately functional entanglement pair in *E. coli* (35) and ported it into the soil microbe, *Pseudomonas protegens Pf-5*. This organism was chosen because it is an attractive rhizophore-dwelling candidate for hosting bioengineered circuits due to its plant-promoting functions and lack of IS elements (52), which are known to contribute to circuit inactivation (13,18,53). The entanglement pair is comprised of a toxin (*relE;* 288 bp) embedded in the +1 frame of a conditionally essential gene (*ilvA;* 1,542 bp). The gene *relE* encodes a mRNA-degrading endoribonuclease of the type II toxin-antitoxin family and *ilvA* encodes threonine deaminase, which is required for isoleucine biosynthesis (54-56). Threonine deaminase is comprised of two primary domains—the N-terminal catalytic domain, which catalyzes the production of isoleucine and the C-terminal regulatory domain, which provides positive and negative conformational feedback to the catalytic domain based on the availability of substrate (threonine and valine) and product (isoleucine) (57,58) (**Fig. S1A**). The gene encoding the toxin *relE* was entangled into the C-terminal domain of *E. coli ilvA*. To accommodate a WT amino acid sequence for RelE, the *ilvA* sequence was significantly recoded with missense mutations (∼79% of entangled residues were altered) (**Fig. S1B**). This pairing allows us to test whether an essential function (isoleucine biosynthesis) can improve the stability of a gene prone to mutational inactivation (*relE*).

Previously, in *E. coli*, the *ilvA*/*relE* entangled pair was found to rescue the growth of an isoleucine auxotroph (Δ*ilvA*) in minimal medium, indicating that this entangled, recoded *ilvA* variant encodes an active enzyme (35). However, *ilvA*/*relE* minimally inhibited cell growth in rich medium, suggesting weak RelE activity (35). To test the functionality of the *ilvA/relE* entanglement construct in *P. protegens*, we first probed the ability of recoded *ilvA* to rescue growth of a strain made auxotrophic for isoleucine (Δ*ilvA*Δ*tdcB*; **Fig. S2**). To test recoded *ilvA* function separately from *relE* toxicity (35), we used an *ilvA*/*relE*^stop^ allele that contains mutations that introduce multiple stop codons within the *relE* reading frame but are silent in the *ilvA* frame (35). A rhamnose-inducible promoter (P_*rhaBAD*_) was used to drive the expression of *ilvA/relE* (**Fig. 1A**). Here, addition of rhamnose stimulates a regulatory cascade in which RhaS activates *rhaSR* expression and then RhaR activates the P_*rhaBAD*_ promoter (59). Consistent with previously reported results in *E. coli, ilvA*/*relE*^*stop*^ rescued the growth of a *P. protegens* Δ*ilvA*Δ*tdcB* strain but required a higher concentration of rhamnose compared to the non-entangled WT *ilvA* control (*ilvA*^WT^), suggesting that the product of entangled *ilvA* is partially functional (**Fig. 1B** and **S3**). We next evaluated the activity of entangled *relE* by testing cell growth with chromosomally encoded *relB* antitoxin. A cumate inducible promoter was used to regulate *relB* expression (P_*cymR*_-*relB*), wherein addition of cumate relieves CymR-mediated repression of the P_*cymR*_ promoter (60-62) (**Fig. 1A**). For the *ilvA*/*relE* strain, *relE* toxin induction failed to inhibit cell growth with or without *relB* antitoxin induction (**Fig. 1C** and **S4**), whereas expression of non-entangled *relE*^*WT*^ severely inhibited cell growth in the absence of *relB* antitoxin (**Fig. 1C** and **S4**). These results suggests that *relE* is not functional in the entanglement design (**Fig. 1C** and **S4**).

We hypothesized that the lack of cellular toxicity by entangled *relE* is the result of poor translation, especially as no apparent ribosome binding site (RBS) could be detected within *ilvA* upstream of the entangled *relE* reading frame (**Fig. S5**) (63). While the algorithm used to generate the entanglement designs (CAMEOS) takes into account protein fitness scoring while satisfying the protein co-encoding constraint, it does not automatically install an internal RBS to ensure translation of the internal gene embedded within the larger gene (**Fig. S5**) (35,64). We therefore sought to enhance RelE expression through *post-hoc* optimization of an internal RBS. Accordingly, we manually designed and engineered five *ilvA/relE*^RBS^ constructs with increasing predicted translation rates (64) upstream of the entangled *relE* start codon (**Fig. S5**). Growth assays revealed that four of the newly designed RBSs including *ilvA/relE*^RBS2^, *ilvA/relE*^RBS3^, *ilvA/relE*^RBS4^ and *ilvA/relE*^RBS5^, successfully recovered RelE activity and inhibited cell growth when the antitoxin was not induced (**Fig. 1C** and **S4**). Only *ilvA/relE*^RBS1^ failed to improve toxicity, which was unsurprising as RBS1 had the lowest predicted translation rate (**Fig. S5**).

Importantly, growth inhibition due to *relE* expression was rescued by induction of the *relB* antitoxin in each construct (**Fig. 1C** and **S4**). This confirms proper RelE activity was achieved through increased translation by the newly designed internal RBSs.

Given that optimizing the RBS strength of entangled *relE* also required nonsynonymous point mutations within *ilvA*, we further tested whether the altered RBSs affect recoded *ilvA* function (**Fig. S5**). Induction of *ilvA/relE*^STOP/RBS1^, *ilvA/relE* ^STOP/RBS2^, *ilvA/relE* ^STOP/RBS4^ and *ilvA/relE* ^STOP/RBS5^ rescued growth in minimal medium to a similar degree as the original *ilvA/relE*^STOP^ entanglement, which suggests that the amino acid changes imparted by the altered RelE RBSs did not impair recoded IlvA function (**Fig. 1B** and **S3**). Surprisingly, *ilvA/relE*^*STOP/*RBS3^ yielded a growth phenotype that closely mirrored the non-entangled *ilvA*^*WT*^; robust growth was observed in minimal medium without the addition of rhamnose. These results suggest that the RBS3 optimization either increased activity or abundance of entangled IlvA (**Fig. 1B-C**). Western blots using functional 3xFLAG tagged constructs showed that the RBS3 modification did not alter recoded IlvA protein abundance (**Fig. S6A-B**), implying an improvement in threonine deaminase enzymatic activity instead (See **Supplemental Discussion**). Overall, these data indicate that post-entanglement modification of the internal RBS improved the expression of the internally entangled gene (*relE*) with a positive impact on recoded IlvA functionality. Since RBS3 improved the toxicity of entangled *relE* while also yielding a growth phenotype nearly identical to *ilvA*^*WT*^, we focused our efforts on the *ilvA/relE*^RBS3^ design in follow-on experiments.

### Entanglement with ilvA Enhances Mutational Robustness of a Toxic Genetic Circuit

Next, we sought to determine whether *ilvA/relE*^RBS3^ can increase the robustness of the embedded toxin to inactivating mutations. Because sequence entanglement imposes constraints on the evolution of both genes in the entangled pair, we expect most mutations that arise in *relE* will also be deleterious to the function of recoded *ilvA*. Thus, we expect that growing the *ilvA*/*relE*^RBS3^ strain under conditions that require *ilvA* function would effectively protect *relE* from accruing inactivating mutations. To test this, we employed a toxin escape assay by growing the *ilvA*/*relE*^RBS3^ strain without selective pressure (rich media, + antitoxin) and then plating on non-permissive (+ toxin, - antitoxin) and permissive (+ toxin, + antitoxin) minimal medium (**Fig. 2A**). In the presence of isoleucine, we found that a strain harboring *ilvA*/*relE*^RBS3^ experienced an escape ratio (colony forming units (CFU) under non-permissive/CFU under permissive conditions) of ∼10^−6^, which is comparable to previous observations when *relE* is expressed in *Pseudomonas* (21) (**Fig. 2B**). However, in the absence of isoleucine, the escape ratio is reduced by ∼7-fold, indicating that entanglement can lower the frequency of the population which accumulates RelE inactivating mutations (**Fig. 2B**). Sequencing the surviving colonies from non-permissive plates revealed that the majority of colonies on plates with isoleucine incurred mutations within the P_*rhaBAD*_ promoter and RhaR/RhaS regulators of *ilvA*/*relE*^RBS3^ or large internal truncations that span across the entanglement region (**Fig. 2C**). However, colonies from plates without isoleucine acquired no mutations in the promoter or regulators of *ilvA*/*relE*^RBS3^ (**Fig. 2C**). Instead, these constructs harbored mutations in either the antitoxin regulator CymR or within the *relE* gene (small deletions) (**Fig. 2C**). These more ‘benign’ mutations likely relieve RelE toxicity by increasing RelB antitoxin expression through reduced CymR activity or by directly ablating the catalytic activity of RelE. No mutations solely affecting IlvA activity were observed from plates without isoleucine, supporting the idea that there was selective pressure to maintain functional IlvA under this condition.

**Fig 2.**
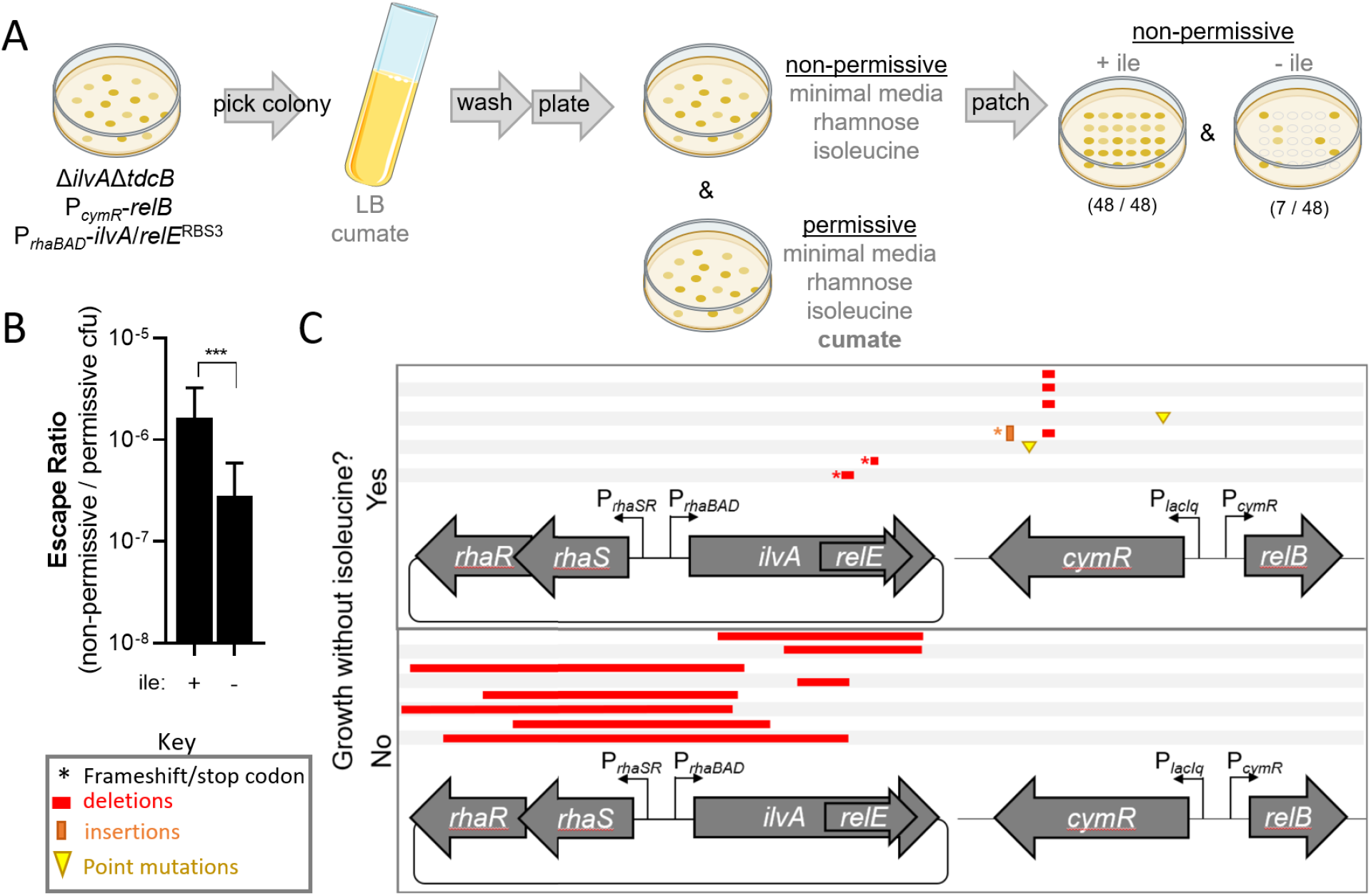
Escape ratio and mutational landscape for *ilvA*/*relE*^RBS3^ following a shift from permissive to non-permissive conditions. (**A**) Diagram of the procedure used to determine the escape ratio. Single isolates of the strain Δ*ilvA*Δ*tdcB* P_*cymR*_-*relB* harboring a vector with *ilvA*/*relE*^RBS3^ were grown under permissive conditions (LB + cumate) and then plated for colony forming units (CFU) on minimal medium under both non-permissive (toxin induced) and permissive (anti-toxin induced) conditions with isoleucine. For each replicate, 48 colonies were then patched from the non-permissive plate with isoleucine onto non-permissive plates either with or without isoleucine. (**B**) The escape ratio was used as a metric of *relE* function and was determined by dividing the CFU/mL on non-permissive medium by the CFU/mL on permissive medium with isoleucine before patching. This was then multiplied by the proportion of colonies that survived on the patched plates either with isoleucine (+ile) or without isoleucine (-ile), respectively. An example of this proportion is seen in **A**. Data are shown as the mean ± SD of 8 independent replicates. Comparisons were made by Student’s T-test. *** = P < 0.001. (**C**) Schematic showing the types of mutations present in toxin escape colonies isolated from non-permissive patch plates without (top) or with isoleucine supplementation (bottom).

The finding that small deletions causing frameshifts within the *relE* entangled region failed to ablate threonine deaminase function raised the question of whether *ilvA* can protect against any mutations within entangled *relE*. To test this, we generated a truncation of *ilvA* just upstream of the entanglement position and the modified internal RBS that removes the entire C-terminus of IlvA (*ilvA*^ΔH322-G514^ (**Fig. S1B**) and assessed whether this construct could support growth in minimal medium. We found that, *ilvA*^ΔH322-G514^ grew similarly to the *ilvA*^*WT*^ strain which suggests that the C-terminal regulatory domain of IlvA is dispensable for function in *P. protegens* (**Fig. S7**). While mutations within *relE* (which is entangled into the C-terminal domain of IlvA) are a potential failure mode of this entanglement design, they are observed less frequently than those occurring within the *ilvA* regulatory elements. As such, these results demonstrate that this entanglement design protects *relE* from the most common inactivating mutations, thereby increasing its mutational robustness and maintaining a low toxin escape ratio.

### Entanglement Enhances Long-Term Evolutionary Stability of a Toxic Circuit

The finding that entanglement with *ilvA* can protect *relE* from certain inactivating mutations suggested that the *ilvA/relE*^*RBS3*^ may preserve a low toxin escape ratio over time and improve kill-switch stability. Previous deployments of *relE* in a kill-switch circuit showed that *relE* is inherently unstable and is rapidly inactivated by mutations during serial passaging even under permissive conditions (i.e., with anti-toxin expression) (21). This is likely due to a growth defect imparted on cells when *relE* is expressed, which is most clearly seen in the absence of the antitoxin (**Fig. 1C** and **Fig. S8**; compare *ilvA*/*relE*^RBS3^ and *ilvA*/*relE*^STOP/RBS3^ without cumate). We hypothesized that 1) mutations that inactivate entangled *relE* function will rapidly outcompete the original design due to fitness burden imposed by RelE toxicity, and 2) growing these lineages in the absence of isoleucine may effectively slowdown the accumulation of *relE*-inactivating mutations within *ilvA*/*relE*^RBS3^. Accordingly, we assessed the stability of the P_*rhaBAD*_-*ilvA*/*relE*^RBS3^ circuit over 100+ generations in medium with or without isoleucine. We conducted serial passaging of 20 independent lineages of Δ*ilvA*Δ*tdcB* P_*cymR*_-*relB* carrying the P_*rhaBAD*_-*ilvA*/*relE*^RBS3^ vector under permissive conditions (+ toxin, + antitoxin). To determine the stability of the construct, each lineage was plated for CFU after each passage on both permissive (+ toxin, + antitoxin) and non-permissive (+ toxin, - antitoxin) medium supplemented with or without isoleucine in accordance with the condition they were passaged in. This escape ratio is used as a proxy for *relE* activity. All *ilvA*/*relE*^RBS3^ lineages grown with isoleucine saw a dramatic increase in escape ratio during the first ∼30 generations (from 10^−6^ to 10^−1^) (**Fig. 3A**). In the following ∼40 generations, the escape ratio remained between 10^−1^ – 10^0^, indicating *relE* gene was rendered non-functional in nearly the entire population (**Fig. 3A**). We also probed the stability of non-entangled *relE*^RBS3^, which is driven by the same RBS3 modification and displayed comparable toxicity to *ilvA*/*relE*^RBS3^ (**Fig. S8**). All lineages of the non-entangled *relE*^*RBS3*^ reached an escape ratio of ≥ 10^−1^ at ∼40 generations (**Fig. S9**). This result confirms that when passaged with isoleucine, kill-switch circuits using *relE*^RBS3^ and *ilvA*/*relE*^RBS3^ are highly unstable.

**Fig 3.**
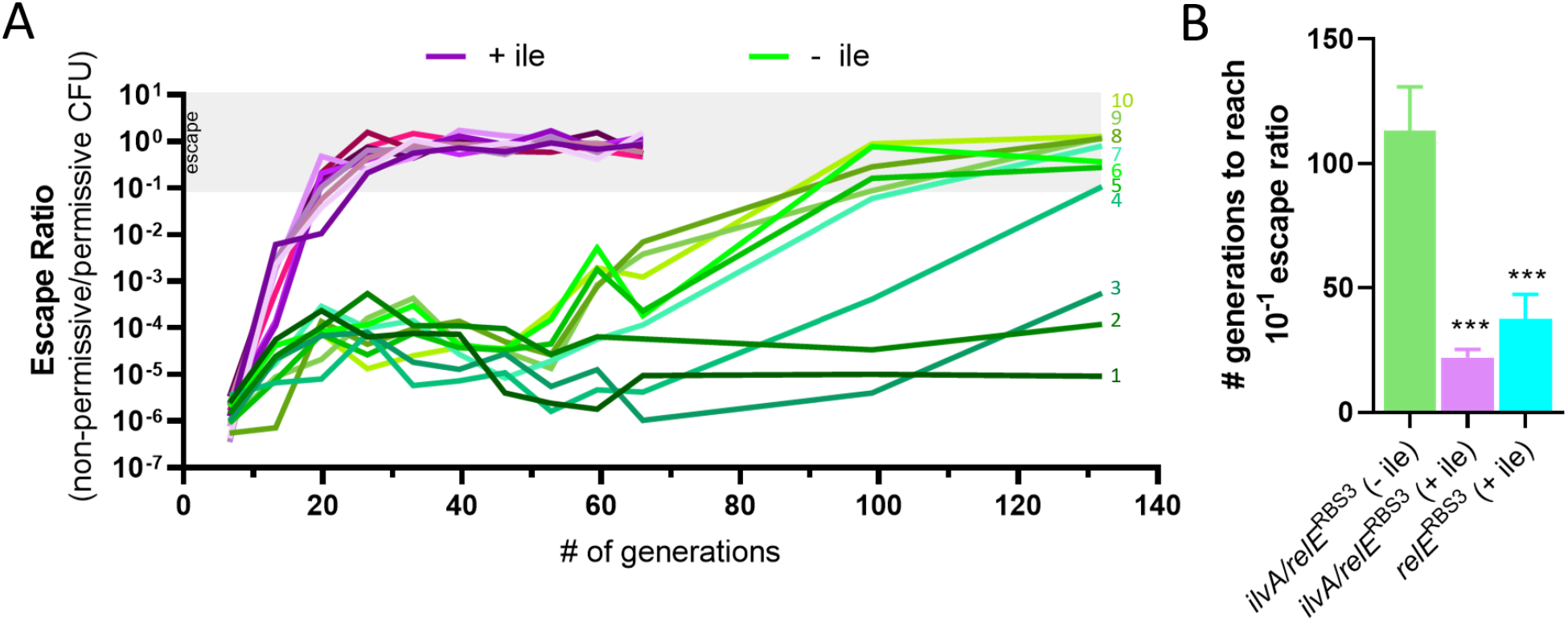
The *ilvA*/*relE*^RBS3^ entanglement increases the evolutionary stability of *relE*. (**A**) Independent lineages of Δ*ilvA*Δ*tdcB* P_*cymR*_-*relB* harboring *ilvA*/*relE*^RBS3^ on a vector were grown in minimal medium with rhamnose (to induce *ilvA*/*relE*^RBS3^) and cumate (to induce antitoxin) in the presence (purple) or absence (green) of isoleucine. Each day (∼6.6 generations) the cultures were diluted 1:1,000 in fresh medium and plated for CFU on toxin permissive and non-permissive conditions and an escape ratio was calculated. After 10 days, measurements were taken less frequently (every 5 days). Passaging in medium with isoleucine was discontinued at ∼66 generations when the population was stabilized with a near 100% escape ratio. Each condition is represented by 10 independent lineages which started from single colonies and are numbered in order of increasing final escape ratio. The grey bar indicates toxin escape ratio ≥ 10^−1^ (10%). (**B**) Average number of generations elapsed when the escape ratio exceeded 10^−1^ (i.e., 10%, grey bar in **A**). Raw data for *relE*^RBS3^ can be found in **Fig. S8**. The result for *ilvA*/*relE*^RBS3^ grown without isoleucine is an underestimate as 3 lineages from this condition never reached 10% escape and were excluded from the calculation. Bar graph data is shown as the mean ± SD. Asterisks directly above bars denote comparisons to the *ilvA*/*relE*^RBS3^ without isoleucine condition (-ile). Comparisons were made by one-way ANOVA with Tukey’s post test. *** = P < 0.001.

In contrast to medium with isoleucine, all *ilvA*/*relE*^RBS3^ lineages passaged without isoleucine (a condition that selects for IlvA function) exhibited a more gradual increase in escape ratio. A slight increase in escape ratio from 10^−6^ to 10^−4^ was observed for all lineages within the first ∼20 generations, before leveling off for the next ∼30 generations (**Fig. 3A**). At ∼50 generations, a divergence in evolutionary trajectory took place. Seven of the lineages experienced a clear upward trend, reaching an escape ratio ≥10^−1^ by ∼113 generations (**Fig. 3A** lineages 4-10; **3B** green bar). However, the escape ratio for three of the lineages passaged without isoleucine remained low (∼10^−5^) even after 132 generations (**Fig. 3A**, lineages 1-3). This marks a significant improvement over the *ilvA*/*relE*^RBS3^ and *relE*^RBS3^ lineages grown with isoleucine, which reached an average escape ratio of 10^−1^ by ∼22 generations and ∼37 generations, respectively (**Fig. 3B**). Together, these data show that *relE* is more genetically stable and less susceptible to inactivating mutations when entangled with *ilvA* and passaged under conditions in which *ilvA* is required for growth.

### Entanglement Alters the Landscape of Allowable Inactivating Mutations

To decipher the genetic basis for circuit stabilization by sequence entanglement, we sequenced the entire P_*rhaBAD*_-*ilvA*/*relE*^RBS3^ plasmid and P_cymR_-*relB* region from one colony randomly selected from the final passage of each lineage grown on a permissive plate (named isolates 1-10, **Fig. 3A**). While we recognize that a single colony does not represent the entire population, it provides an indication of the types of mutations selected for under these growth conditions. For all lineages grown with isoleucine, large deletions were observed in the plasmid spanning both the *rhaS* and/or *rhaR* regulatory genes and the *ilvA*/*relE*^RBS3^ entanglement region which undoubtedly disrupted the function of both the toxin and recoded IlvA (**Fig. 4A**). This mutation pattern is similar to our findings from the single time-point escape assay in which escape mutants were sequenced from medium containing isoleucine (**Fig. 2C**). In contrast, such large and disruptive deletions were not observed in any of the lineages grown without isoleucine, indicating that the selective pressure to maintain IlvA function alters the landscape of allowable inactivating mutations in the circuit (**Fig. 4A**). The colonies from the three lineages with the highest final escape ratio in the absence of isoleucine (**Fig. 3A**, isolates 8, 9, 10) contained a deletion of the entire *relE* gene (**Fig. 4A**), which is an expected failure mode for this entanglement given the dispensability of the C-terminal portion of IlvA (**Fig. S7**). The other isolates from lineages grown without isoleucine contained point mutations or smaller insertions and deletions in the genes encoding the regulators *rhaR* and *rhaS* (**Fig. 4A**). None of the isolates contained a mutation in the chromosomal P_cymR_-*relB* region (**Fig. 4A**). We posit that these lineages, derived through adaptive evolution, led to improved circuit expression levels by reducing the burden of RelE while maintaining sufficient IlvA production. Therefore, we examined the vectors from these isolates to determine how the mutations may impact *ilvA*/*relE* activity.

**Fig 4.**
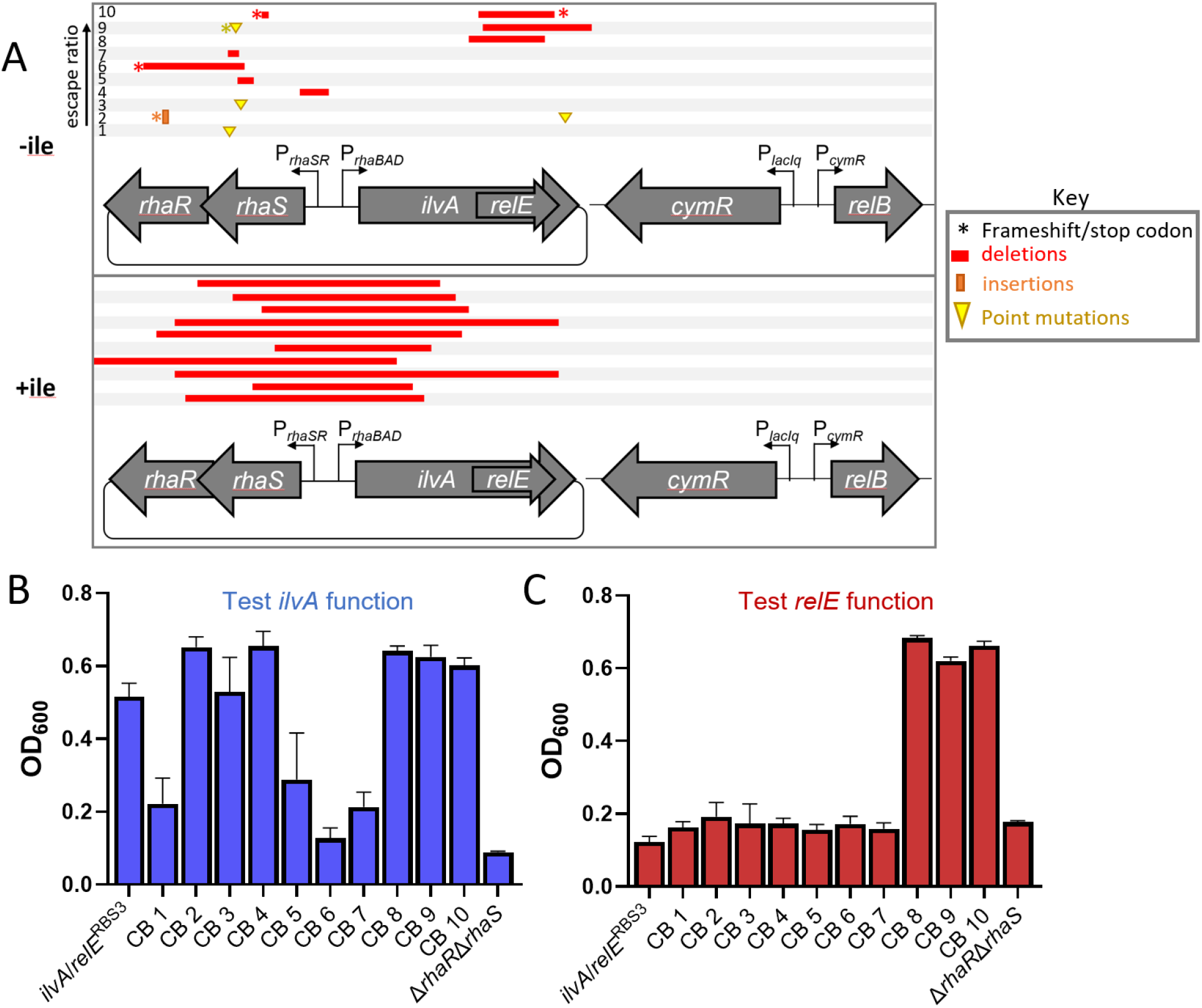
The *ilvA*/*relE*^RBS3^ entanglement alters the mutational landscape and maintains function of both genes. (**A**) Schematic of the types of mutations present in colonies isolated from each lineage of the long-term evolutionary stability assay either grown with or without isoleucine. A single colony was isolated from each lineage under permissive conditions after the final passage. Mutations were identified by sequencing the entire vector and the *P*_*cymR*_*-relB* chromosomal region. Sequencing results from each isolate are listed in order of increasing final escape ratio seen in **Fig. 3A** (isolates 1-10). The isolated vectors were transformed into a clean genetic background (Δ*ilvAΔtdcB P*_*cymR*_*-relB)* and were grown in (**B**) minimal medium without isoleucine and with addition of rhamnose and cumate to probe *ilvA* function or (**C**) in minimal medium with isoleucine and rhamnose and without cumate to test *relE* toxicity. For comparison with these clean background (CB) strains, the original parent strain (*ilvA*/*relE*^RBS3^) and a strain in which the regulators *rhaR* and *rhaS* are deleted from the vector (Δ*rhaR*/*rhaS*) are included. Growth is reported as OD_600_ after 15 h. Data are shown as the mean ± SD of 3 independent replicates.

We first assessed the toxicity of entangled *relE* and the function of recoded *ilvA* for the colonies isolated from the final passage of each lineage grown without isoleucine (isolates 1-10). When grown in minimal medium in permissive conditions (+ toxin, + antitoxin) without isoleucine, each isolate exhibited a shorter lag phase and a similar density after 15 hours of growth compared to the parent strain (**Fig. S10A-B**), indicative of a functional IlvA. Next, we assessed entangled *relE* function by growing the strains in minimal medium under non-permissive conditions (+ toxin, - antitoxin) supplemented with isoleucine. We found that colonies isolated from lineages 8, 9 and 10 grew unimpeded, indicating a loss of *relE* activity (**Fig. S10C-D**). In contrast, isolates 1-7 exhibited a significant growth defect, indicative of functional *relE* (**Fig. S10C-D**). This result demonstrates that *relE* toxicity was preserved in at least a subset of the population for lineages 1-7 (**Fig. 3A**).

To ascertain the role of the mutations in the evolved *ilvA*/*relE*^RBS3^ vectors in preserving fitness and maintaining or diminishing toxicity, we purified plasmids from the isolated colony of each lineage grown without isoleucine (i.e., from isolates 1-10) and transformed them into a <u>c</u>lean genetic <u>b</u>ackground (*P. protegens* Δ*ilvA*Δ*tdcB* P_*cymR*_-*relB*) (named CB strains 1-10). We then independently assessed the isoleucine auxotrophy and *relE* toxicity of these new CB strains, as described above. These data showed that CB 8, 9 and 10 supported growth in both permissive and non-permissive conditions, which confirmed that the deletion mutations in these vectors inactivated RelE while maintaining a functional IlvA (**Fig. 4B-C**). Similar to the original isolates, CB 1-7 also displayed diminished growth under non-permissive conditions, which suggests these vectors all contained functional *relE* (**Fig. 4C**). While there was a delay in the growth inhibitory effect of *relE* compared to the parent strain (**Fig. S11A**), the final OD_600_ for all CB strains with vectors from isolates 1-7 was similar to the parent *ilvA*/*relE*^RBS3^ strain (**Fig. 4C** and **S11A**). This result demonstrates that the mutations in the *rhaR*/*rhaS* regulator region of CB 1-7 still permitted sufficient entangled RelE expression.

To probe the function of recoded IlvA in these vectors, we grew CB strains 1-7 in permissive conditions in the absence of isoleucine, which revealed that all seven vectors with functional *relE* supported some growth in the clean background, although there were notable differences in the lag phase of growth (**Fig. S11B**). CB strains 1, 5, 6, and 7 appeared more compromised in their ability to make isoleucine as evidenced by an extended lag phase and lower OD_600_ after 15 hours of growth compared to the parent strain (**Fig. 4B** and **S11B**). This is in contrast with the results from the original colony isolates which showed all strains can grow in minimal medium under permissive conditions (**Fig. S10A-B**). Thus, we suspect that there may be compensatory mutations elsewhere in the chromosome that contribute to isoleucine biogenesis in the original colonies isolated from lineages 1, 5, 6 and 7. However, CB 2, 3 and 4 yielded robust growth compared to the parent construct, and CB strains 2 and 4 supported a shorter lag phase (**Fig. 4B** and **S11B**). The mutations in CB 2, 3 and 4 include a 1bp insertion in *rhaR* (*rhaR*^L128+1bp^), a missense point mutation in *rhaS* (*rhaS*^W190G^), and a deletion of the entire P_*rhaSR*_ promoter including the first 15bp of *rhaS* (*rhaS*^ΔM1-H5^/P_*rhaSR*_^Δ179bp^), respectively. With these mutations, it is possible that RhaS is still partially functional and can activate P_*rhaBAD*_. In contrast, CB strains 1, 5, 6 and 7 contain vectors with mutations which likely impact RhaS activity and more severely affect the expression of *ilvA*/*relE*^RBS3^: CB 1 has a substitution in the *rhaS* DNA binding domain (*rhaS*^F254L^) that has been shown to be critical for RhaS activity (65) and CB 5, 6 and 7 harbor deletions within the DNA binding domain of RhaS (**Fig. 4A**). CB 2 also accumulated a point mutation in the IlvA C-terminus (*ilvA*^G455C^) of the *ilvA*/*relE*^RBS3^ construct downstream of entangled *relE*; however, since the C-terminus of IlvA is not required for isoleucine biogenesis (**Fig. S7**), this mutation does not likely impact IlvA or RelE function. Due to the growth phenotypes observed among these CB strains, we hypothesize the mutations in CB 2, 3 and 4 reduce the strength of the P_*rhaBAD*_-*ilvA*/*relE*^RBS3^ expression to a level compatible with functional IlvA production and reduced RelE burden under permissive conditions, while still preserving RelE toxicity under non-permissive conditions.

### Sequence Entanglement Allows for Evolution to Optimize Circuit Expression Which Improves Fitness and Genetic Stability

To determine if the mutations in CB 2, 3 and 4 improve cellular fitness compared to the original parent strain, we employed a competitive growth assay (66). We hypothesized that if the vectors from isolates 2, 3 and 4 imparted a competitive growth advantage to their respective host strains, then they should outcompete the parent strain if grown together. Indeed, a competitive growth assay, in which the original P_*rhaBAD*_-*ilvA*/*relE*^RBS3^ strain was grown in co-culture with CB 2, 3 or 4, revealed a competitive index between ∼37- and ∼51-fold greater than the parent strain after 13 generations of growth in permissive conditions (**Fig. 5A**). These data reveal that the mutations in the P_*rhaBAD*_ promoter of CB 2, 3, and 4 likely optimize the expression of their respective *ilvA*/*relE*^RBS3^ circuit and impart a significant growth advantage compared to the parent strain.

**Fig 5.**
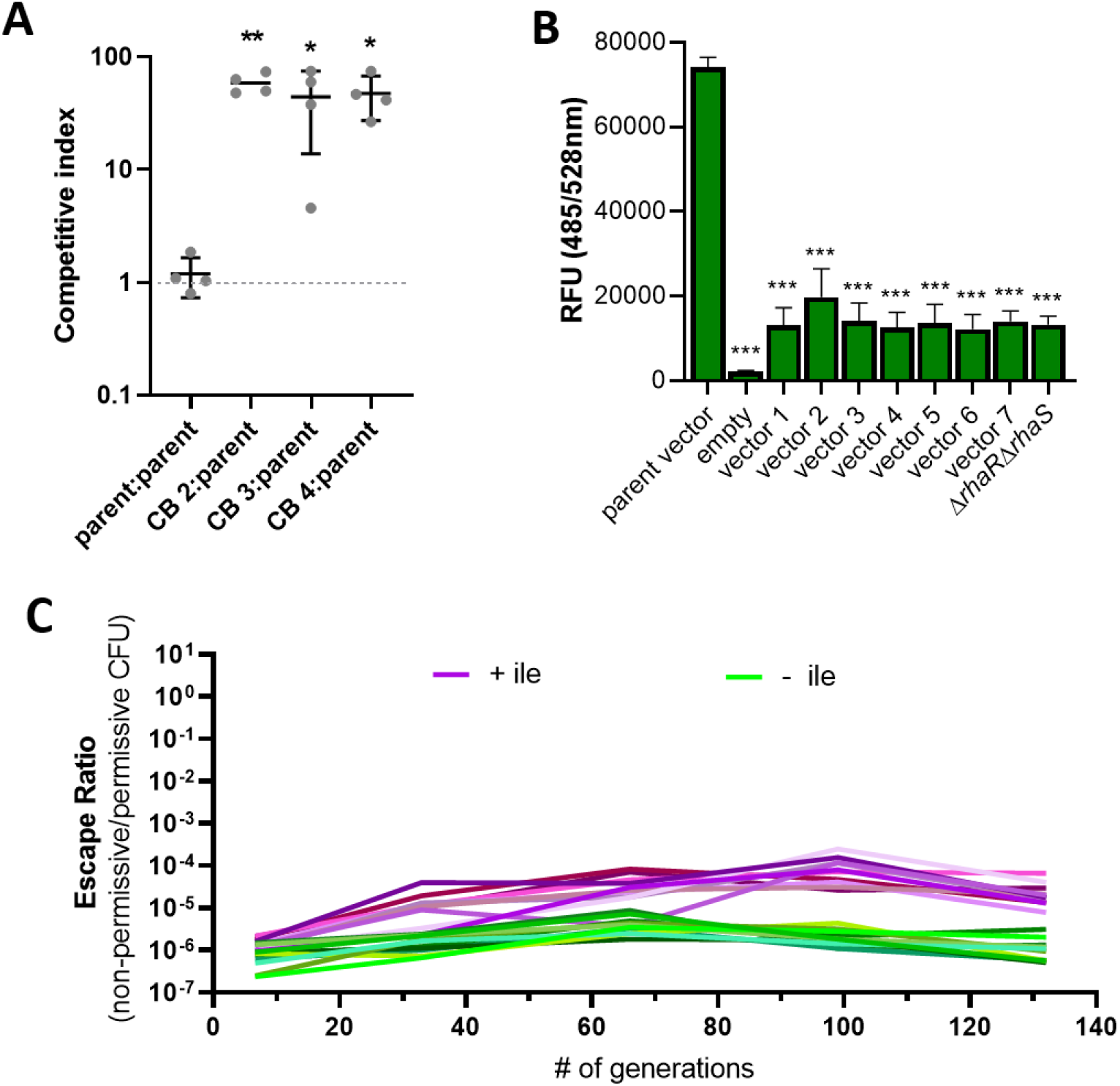
Sequence entanglement allows for circuit optimization through adaptive evolution. (**A**) Competition assay in which the parent strain *ilvA*/*relE*^RBS3^ was grown in a 1:1 co-culture with clean background strains harboring vectors isolated from the listed lineages (CB 2-4). Strains were differently marked with chromosomally integrated antibiotic cassettes, tetracycline and gentamicin. The competitive index was calculated as the CFU ratio of mutant/parent after growth for 48 hours divided by the CFU ratio of the mutant/parent in the initial inoculum. (**B**) Assay of promoter activity for vectors isolated from lineages 1-7 of the long-term evolutionary stability assay. The *gfp gene* was cloned downstream of the promoter of each vector and strains were grown in minimal medium overnight with rhamnose. RFU is reported relative to OD_600_. (**C**) Independent lineages of CB 2—a clean genetic background (Δ*ilvA*Δ*tdcB* P_*cymR*_-*relB*) strain harboring the *ilvA*/*relE*^RBS3^ vector isolated from the original lineage 2 in **Fig. 3B**—were grown in minimal medium with rhamnose (to induce *ilvA*/*relE*^RBS3^) and cumate (to induce antitoxin) in the presence (purple) or absence (green) of isoleucine. Every 5 days (∼33 generations) the cultures were diluted 1:1,000 in fresh medium and plated for CFU under toxin permissive and non-permissive conditions and an escape ratio was calculated. Each condition is represented by 10 independent lineages which started from single colonies grown on permissive medium. Data for panels **A** and **B** are shown as the mean ± SD of 3 to 4 independent replicates. Asterisk(s) directly above data denote comparisons to the parent condition. Comparisons were made by one-way ANOVA with Dunnet’s (**A**) or Tukey’s (**B**) post test. *** = P < 0.001, ** = P < 0.01, * = P < 0.05.

We next sought to understand how the accumulated mutations may affect circuit expression and by what mechanism these mutations are improving fitness. Given the delay in the onset of *relE* toxicity with the vectors from isolates 1-7 (**Fig. S11A**) and the fact that each vector had mutations in *rhaR*/*rhaS* (**Fig. 4A**), it seemed likely that these mutations reduced P_*rhaBAD*_-*ilvA*/*relE*^RBS3^ expression. To test this, we replaced *ilvA*/*relE*^RBS3^ with *gfp* in the plasmids from isolates 1-7 (named vectors 1-7) and evaluated GFP expression as a readout for promoter strength. Since many of these vectors contained mutations in *rhaR* and *rhaS* that we might expect to ablate activity of these regulators (i.e., deletions, frameshifts, and point mutations), we also assessed the activity of a vector in which *rhaR* and *rhaS* were deleted (Δ*rhaR*/*rhaS*) for comparison. Interestingly, this analysis demonstrated that all seven vectors exhibited lower fluorescence compared to the original parent vector but were indistinguishable from the Δ*rhaR*Δ*rhaS* vector control (**Fig. 5B**). We expected that GFP induction might be slightly higher in vectors 2, 3 and 4 because the mutations in these constructs still allowed for robust growth in minimal medium in a clean genetic background (**Fig. 4B**), in contrast to a strain harboring a Δ*rhaR*Δ*rhaS* vector which did not support growth; however, no statistical difference in fluorescence was observed between vectors 1-7 (**Fig. 4B** and **5B**). It is possible this *gfp* assay was not as sensitive as the growth-based assays in distinguishing between the low levels of induction observed in these vectors. Nevertheless, these data suggest that the mutations within the RhaR and RhaS regulators of each evolved lineage reduce P_*rhaBAD*_-*ilvA*/*relE*^RBS3^ expression, and this reduced expression is likely responsible for the improved fitness observed in CB strains 2, 3 and 4 (**Fig. 5A**). This result is consistent with previous findings which recommend lowering the expression of a toxic circuit for maintaining cellular fitness and circuit function (14,20,31,67,68).

We thus considered the intriguing possibility that mutations found in CB 2, 3 and 4 have decreased the pressure to select for mutations that abolish entangled *relE* toxicity, thereby enhancing the mutational stability of the toxin *relE (69)*. To test this, CB 2 was chosen for passage over the course of 100+ generations under the same conditions as the initial long-term stability experiment in **Fig. 3A**. Remarkably, all the CB 2 lineages grown under conditions with or without isoleucine exhibited a low, stable escape ratio, never reaching higher than 10^-4 (**Fig. 5C**). This is in sharp contrast to the rapid escape observed with the original parent strain during growth with isoleucine (**Fig. 3A**). The CB 2 lineages passaged without isoleucine maintained an escape ratio between ∼10^−7^–10^−5^ over ∼132 generations. When passaged with isoleucine, CB 2 lineages are also quite stable and displayed only a ∼50-fold increase in escape ratio over the same time frame (**Fig. 5C**). Together, these results suggest that the vector from the original isolate 2 (with the *rhaR*^L128+1bp^ insertion) imparts greatly enhanced *relE* long-term stability compared to the original parent vector (**Fig. 3A** and **5C**). This implies that even under conditions in which *ilvA* is not required for growth, the basal toxicity of *ilvA*/*relE*^RBS3^ from CB 2 does not impart a meaningful fitness defect under permissive conditions. Therefore, mutations that lowered the expression of *ilvA*/*relE*^*RBS3*^ appear to strike a balance between lessening *relE* toxicity while still maintaining *ilvA* function. In conclusion, these data demonstrate that sequence entanglement paired with adaptive evolution can optimize circuit function for prolonged evolutionary stability.

## DISCUSSION

When there is no selective pressure to maintain the function of a burdensome genetic circuit, loss-of-function mutations tend to accumulate in the circuit and cells with broken circuits overtake the population (70,71). The development of mechanisms to stabilize genetic circuits is an important step in deploying engineered biological systems for human benefit. The data presented here shed light on the utility of sequence entanglement to increase the long-term evolutionary stability of costly gene circuits in an environmentally relevant microbe. Below, we discuss key findings from the model *ilvA*/*relE* entanglement pair examined in this study.

Despite having *relE* entangled within a dispensable region of *ilvA* (the C-terminal regulatory domain), the *ilvA/relE*^*RBS3*^ entanglement was capable of stabilizing the toxin RelE. While this entanglement arrangement was vulnerable to certain mutations (e.g., frameshifts and deletion within *relE*), it protected against the most common circuit inactivating mutations (i.e., large deletions across the P_*rhaBAD*_ promoter and *ilvA/relE*). As such, this result is conceptually similar to that achieved with the Riboverlap method (37), where a partial gene overlap is created by inserting a translation initiation site for an essential gene within an upstream GOI, which achieves GOI protection against a subset of the mutational spectrum (e.g., frameshifts within the overlap region and promoter inactivating mutations). An *in-silico* analysis predicted that this Riboverlap approach would increase the temporal stability of the GOI under conditions in which the protected class of mutations are the most frequent inactivating mutations, but this result was not tested experimentally. In this study, we empirically demonstrate that entanglement re-directs evolutionary pressure by selecting against high-frequency mutations that inactivate toxin function, which allows for the concomitant optimization of strain fitness and circuit stability during the course of an evolutionary stability assay. Ultimately, this pairing of gene entanglement with adaptive evolution yielded a kill-switch circuit in which all lineages remained functionally stable for >130 generations, which compares favorably with the most stable kill-switch circuits developed to date (72). As such, we posit that an adaptive laboratory evolution campaign following sequence entanglement can be an effective strategy for improving the fitness and stability of synthetic genetic circuits in an unbiased fashion, thereby obviating the need for manual circuit optimization efforts (20).

The concept of linking a burdensome GOI with an essential function to enhance mutational robustness has been previously implemented using non-entanglement based methods such as transcriptionally linking an antibiotic resistance gene to circuit expression (31), using bi-directional promoters to control expression of both an essential gene and GOI (30), and through synthetic addiction using ligand-responsive molecular biosensors that link production of a target metabolite to expression of an essential function (25). While the first two techniques enhanced the evolutionary stability of a circuit to varying degrees, these strategies remain prone to circuit inactivation by large insertions and deletions. Moreover, although synthetic addiction achieved exemplary stability, the availability of biosensor modules limits the broad utility of this strategy. Here we show that sequence entanglement is an alternative and effective strategy for protecting synthetic circuits by directly linking the sequence that confers the desired circuit function to cell viability. Importantly, sequence entanglement does not introduce extra circuit components to the system, minimizing the number of potential breakage points.

An important design consideration for imparting mutational robustness is the choice of the essential gene. Here, *ilvA* was made conditionally essential by knocking out the two annotated paths for isoleucine biosynthesis in *P. protegens* (Δ*ilvA*Δ*tdcB*). One potential limitation of this system could be the accumulation of compensatory mutations that abolish the necessity of *ilvA*. A recent study showed that *E. coli* acquires mutations to circumvent isoleucine auxotrophy at a high frequency in a Δ*ilvA* strain via activation of an underground isoleucine biosynthesis pathway (through *metABC*) (73). Importantly, the *metABC* genes are not present in *P. protegens* and our Δ*ilvA*Δ*tdcB* strain exhibits a low escape ratio in minimal medium (**Fig. S12;** ∼10^−9^), indicating this host cannot easily circumvent isoleucine auxotrophy if not selected for. Additionally, cross-feeding by non-deficient neighbors is another vulnerability. Studies in *E. coli* have shown that an isoleucine auxotroph can be stably maintained in a population containing isoleucine producing variants (73-75). However, population modeling predicts a crash in population size when isoleucine concentration dips below a critical size (74). As such, non-isoleucine producing cheaters likely cannot completely thwart the protective effects of entanglement but may impact population dynamics. Ultimately, the choice of an (conditionally) essential gene for entanglement should not only consider the flexibility and mutational protectiveness, but also be informed by knowledge of the target environment and host-specific modes of mutational escape.

Our results identify key strengths and limitations of the *ilvA*/*relE* entanglement pair that provide important design considerations for the utility of sequence entanglement. Future studies will focus on testing the generalizability of these results using additional entanglement pairs and by targeting entanglement locations within or in proximity to more critical protein domains (e.g., enzyme active site). Ultimately, this represents a successful showcase of entanglement as a strategy to protect and improve the genetic stability of synthetic circuits and may be useful to a wide variety of bioengineering applications. This may be especially relevant for current biocontainment strategies such as kill-switches, auxotrophies, codon recoding and others that are vulnerable to genetic mutations (17-19,76,77).

## ACKNOWLEDGEMENTS

This work is supported by the U.S. Department of Energy (DOE), Office of Science, Office of Biological and Environmental Research, Lawrence Livermore National Laboratory SFA “From Sequence to Cell to Population: Secure and Robust Biosystems Design for Environmental Microorganisms.” This work was performed under the auspices of the U.S. Department of Energy by Lawrence Livermore National Laboratory under Contract DEAC52-07NA27344 (LLNL-JRNL-845594).

## Supplemental Information

## SUPPLEMENTAL DISCUSSION

Based on our observations in this work, we anticipate that entanglement designs will, in most cases, require post-entanglement optimization of an internal RBS to facilitate translation of the internal gene. In this proof-of-concept example, we achieved toxic levels of RelE using a *post*-*hoc* RBS optimization approach whereby strong RBS sites that minimize disruptive amino acid substitutions within *ilvA* were identified and introduced. Fortuitously, one such RBS site (RBS3) not only improved translation from the embedded *relE* gene but also enhanced the functionality of IlvA (**Fig. 1B-C** and **S3**). We posit that the entanglement of WT *relE* within the genetic region encoding the C-terminal regulatory domain of *ilvA* reduces or alters the autoregulation of IlvA; this phenomenon may also explain the diminished functionality of *ilvA*/*relE*^*stop*^ compared to *ilvA*^*WT*^ in minimal medium without isoleucine (**Fig. 1B** and **S1**). Since the RBS modifications occur in the alpha-helical “neck” region that separates the catalytic and regulatory domain (**Fig. S1A**), we speculate that the amino acid changes imposed by RBS3 (i.e. *ilvA*^L333V/G334L^) may have locked the N-terminus of IlvA into a catalytically active state. This would allow for the constitutive production of isoleucine and support the enhanced growth that we observed with the RBS3 modification without altering IlvA expression levels (**Fig. S6B**). Confirming this hypothesis will require further biochemical and structural characterization. While we do not expect this fortuitous result to generalize to other entanglement pairs, it does highlight an important design constraint of entanglement: mutations that improve the strength of the internal RBS may impact the fitness of the external entangled gene. Entanglement algorithms that incorporate RBS modifications along with fitness prediction would be highly valuable for future entanglement designs.

**Fig S1.**
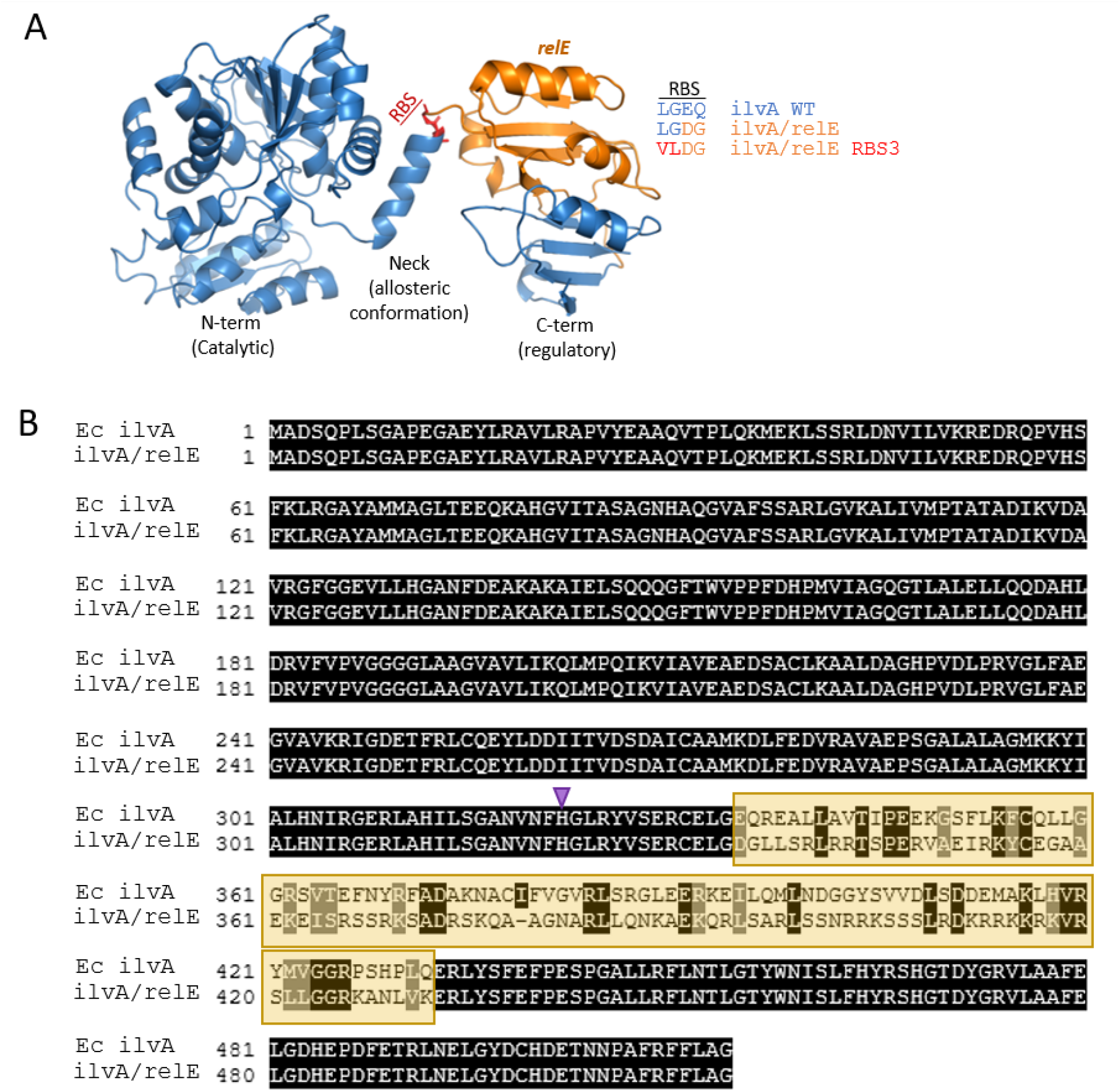
Sequence and structure of *ilvA*/*relE* entanglement. **(A)** Structure of *E. coli* threonine deaminase (blue) with the region of *relE* entanglement highlighted in orange. The amino acid changes imparted by the RBS3 modification in *ilvA*/*relE*^RBS3^ are shown in red. The Protein Data Bank (PDB) accession number is 1TDJ. Structural renderings were generated using PyMOL v. 2.3.4. **(B)** Alignment of the amino acid sequence between *E. coli ilvA*^*WT*^ and the *ilvA*/*relE* entanglement design. The orange box denotes the entanglement position. The purple triangle indicates the position of the C-terminal truncation (*ilvA*^ΔH322-G514^) seen in **Fig. S8**.

**Fig S2.**
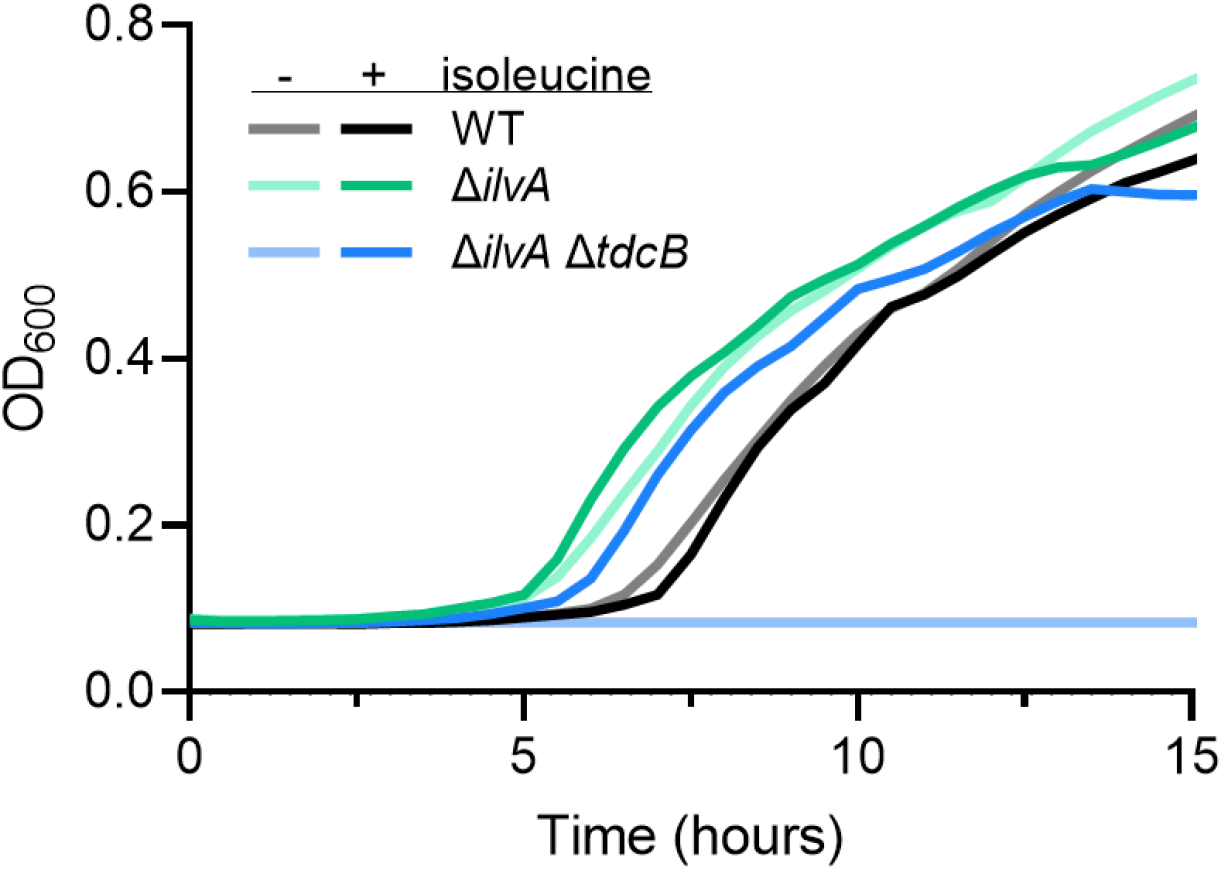
*ilvA* and *tdcB* are required for isoleucine biosynthesis in *P. protegens* Pf-5. In *P. protegens* Pf-5, *ilvA* and *tdcB*, a close homolog of *ilvA*, are both required for isoleucine biosynthesis. Strains were grown in minimal medium with or without the addition of isoleucine. Growth is reported as OD_600_ over time. Data are shown as the mean of 3 independent replicates.

**Fig S3.**
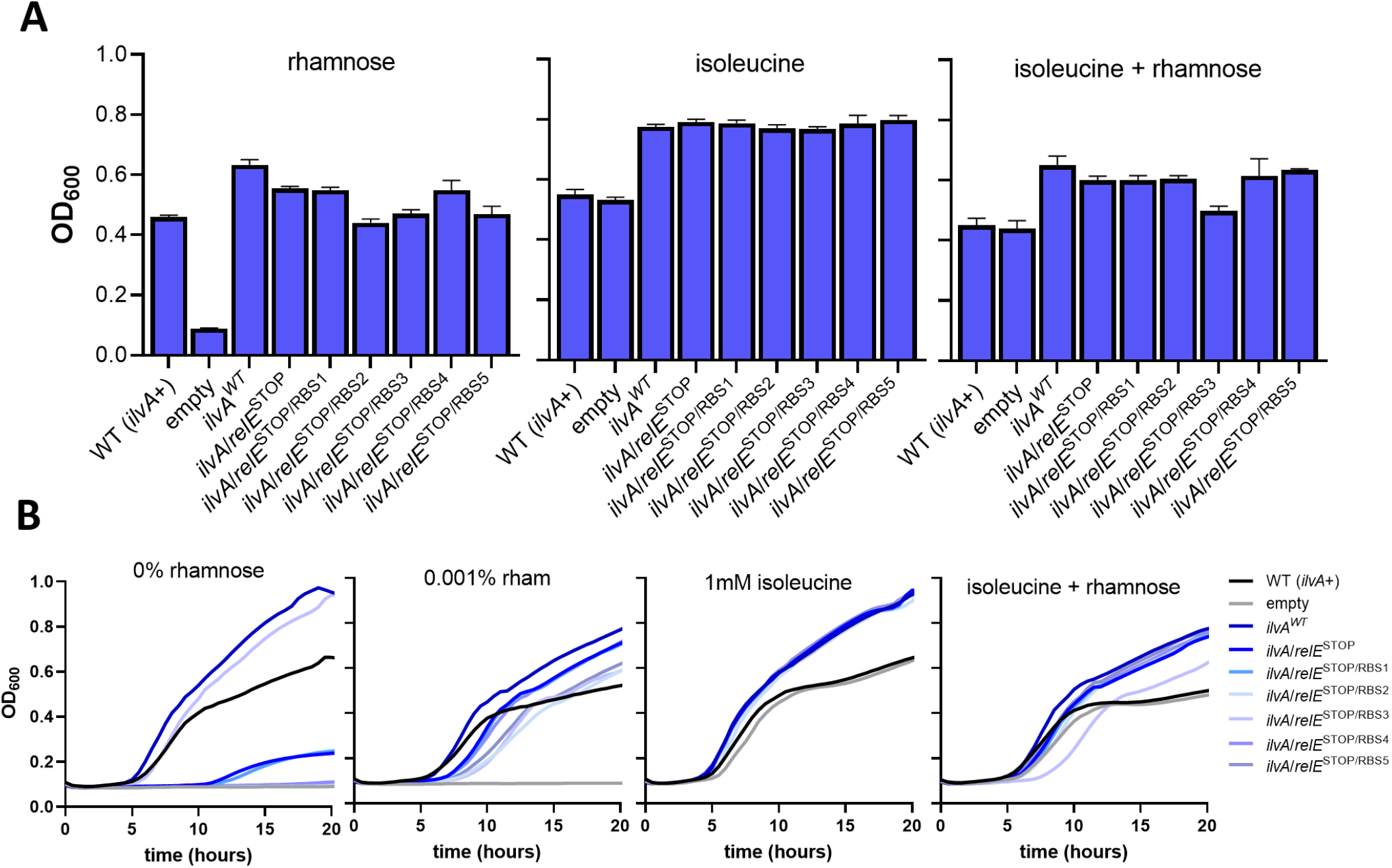
Testing the ability of *ilvA*/*relE*^*STOP*^ alleles to rescue isoleucine auxotrophy. Final density of isoleucine auxotroph (Δ*ilvAΔtdcB)* strains with chromosomally integrated antitoxin (P_*cymR*_-*relB*) harboring plasmids carrying the different alleles listed driven under the P_*rhaBAD*_ promoter. **(A)** Strains harboring *ilvA*/*relE*^*STOP*^ alleles containing different strength RBSs were grown in minimal medium with addition of either rhamnose (to induce the P_*rhaBAD*_ -*ilvA*/*relE* alleles) or isoleucine. Growth is reported as OD_600_ after 15 h. **(B)** Growth curves from data represented in figure **1B** and **S3A** in which growth is reported as OD_600_ over time. All data in bar graphs are shown as the mean ± SD and data in growth curves are shown as the mean. All data are representative of 3 independent replicates.

**Fig S4.**
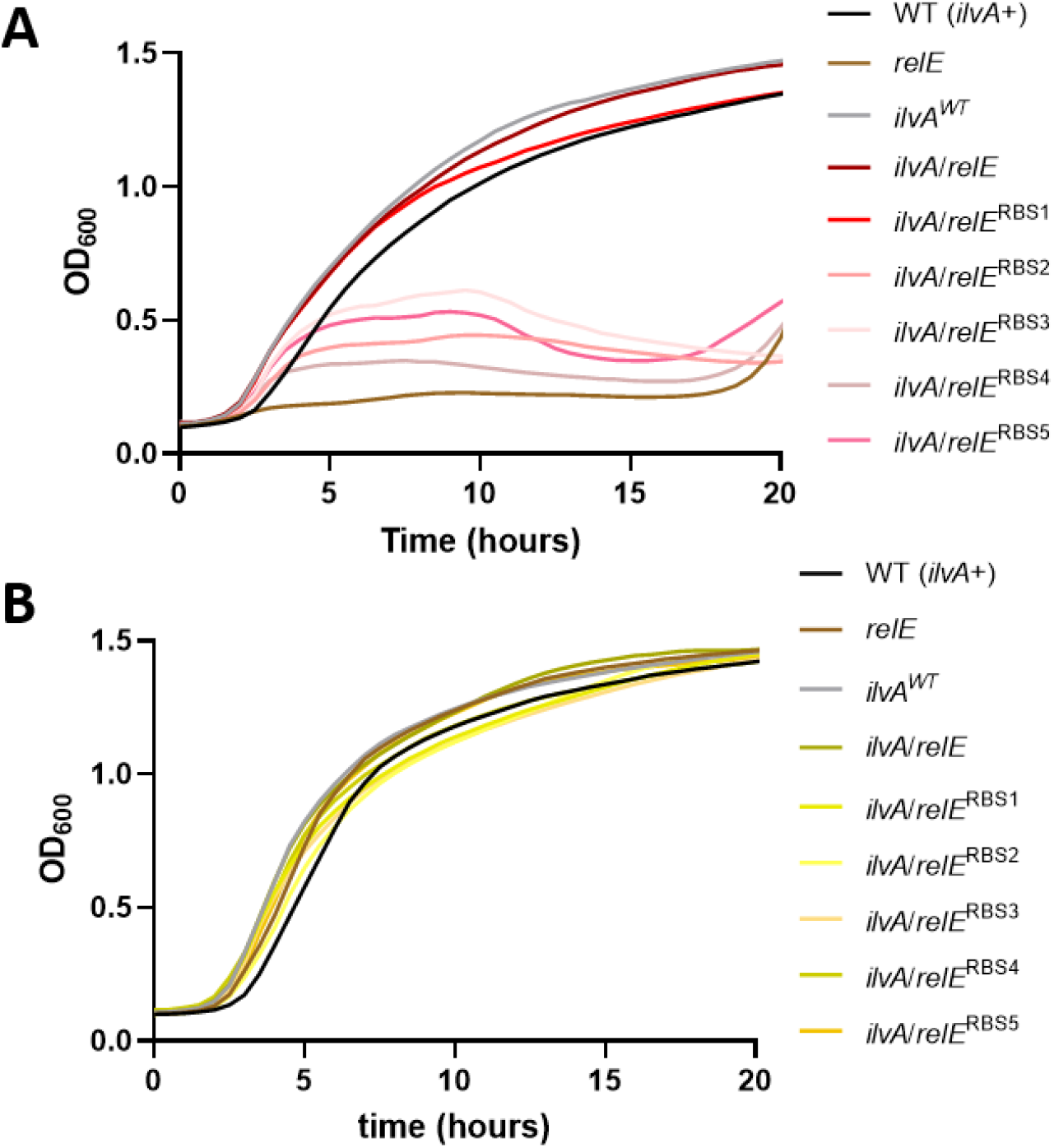
Testing capability of *relE* in *ilvA*/*relE* alleles to suppress growth. Growth curves represent the data from **Fig. 1C** in which growth is reported as OD_600_ over time. (**A**) Strains (Δ*ilvAΔtdcB*, P_*cymR*_-*relB*) harboring P_*rhaBAD*_*-ilvA*/*relE* alleles with different strength RBSs were grown in rich medium to isolate *relE* toxicity. Experiments were performed without rhamnose induction of *ilvA*/*relE* expression (**B**) To rescue growth, the antitoxin was induced by addition of cumate. Data is shown as the mean of 3 independent replicates.

**Fig S5.**
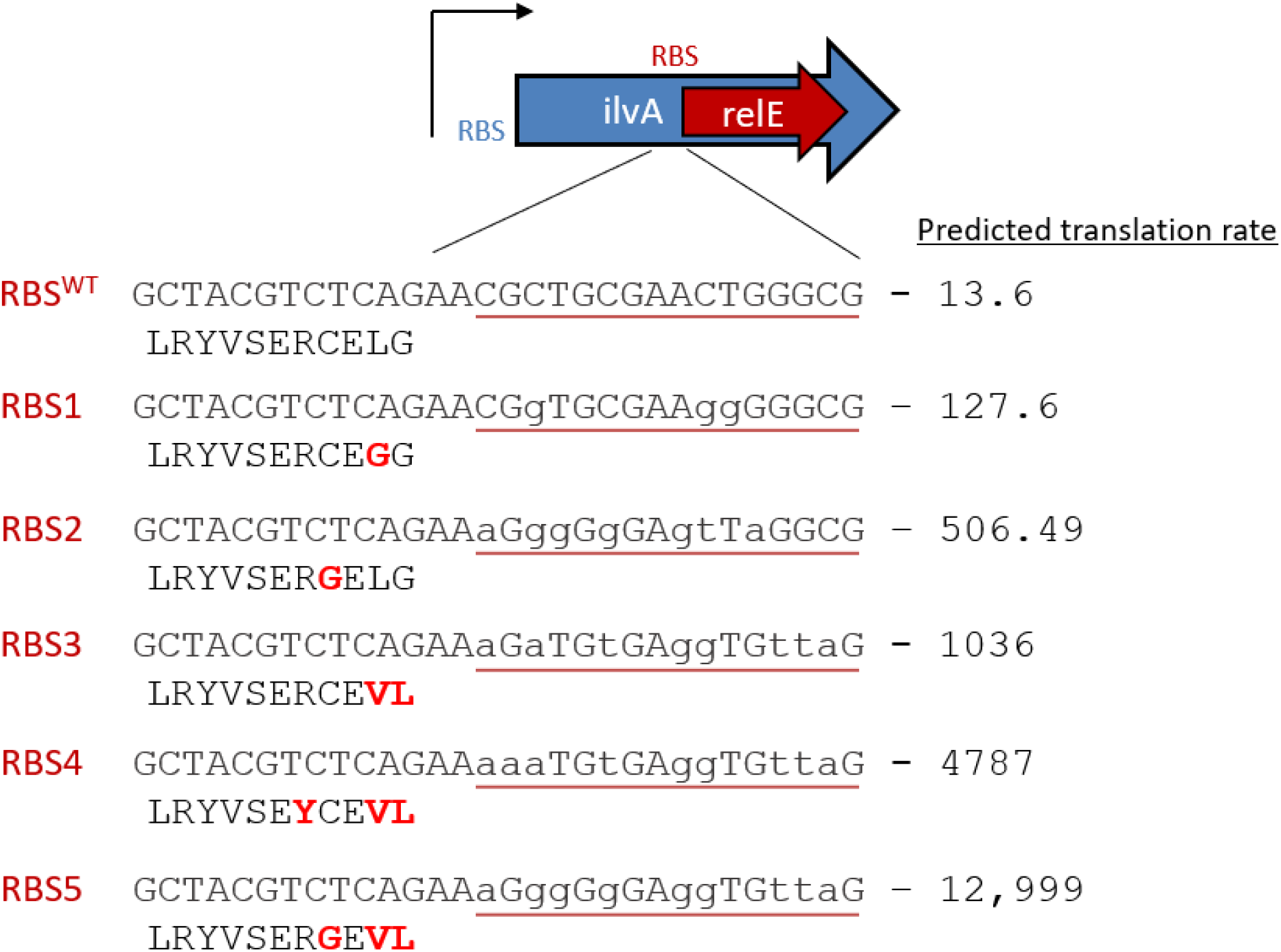
Optimization of internal RBS to improve RelE expression. The nucleotide sequence (top) and amino acid sequence (bottom) of the internal RBS region within *ilvA* (red underline), just upstream of the start codon of entangled *relE*, is listed next to each variant tested. Changes to the nucleotide sequence of *ilvA* are seen with lowercase letters and the corresponding changes in the amino acid sequence are depicted in bold red font. The predicted translation rate was determined using the Salis Lab RBS calculator (1). The rates are given on a proportional scale ranging from 1 to 100,000+.

**Fig S6.**
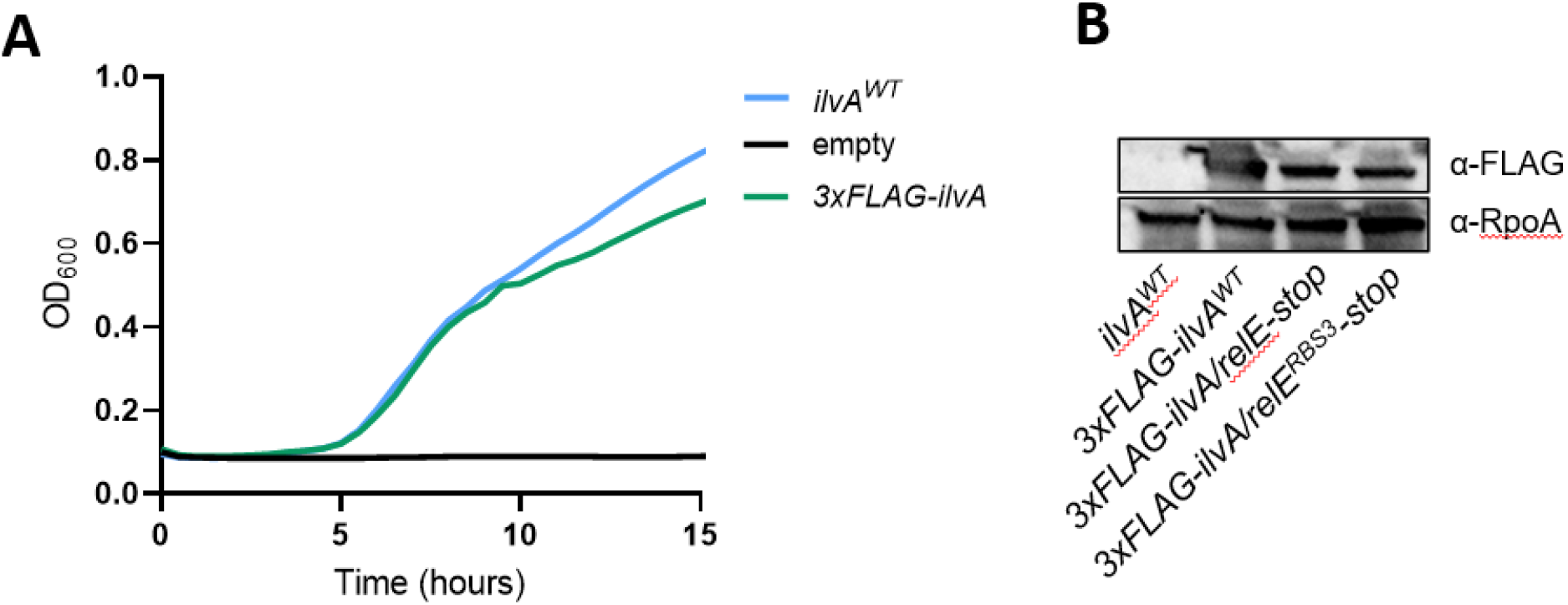
Improved RBS modification does not affect threonine deaminase expression. **(A)** Growth curve in minimal medium without isoleucine shows that 3xFLAG tagged *ilvA* constructs are functional for isoleucine biosynthesis. Growth is reported as OD_600_ over time. **(B)** Western blot testing whether entanglement and/or RBS3 modification affects expression of threonine deaminase. Strains were grown overnight with rhamnose for induction of the *ilvA* variants. Growth curve data are shown as the mean and all data are representative of 3 independent replicates.

**Fig S7.**
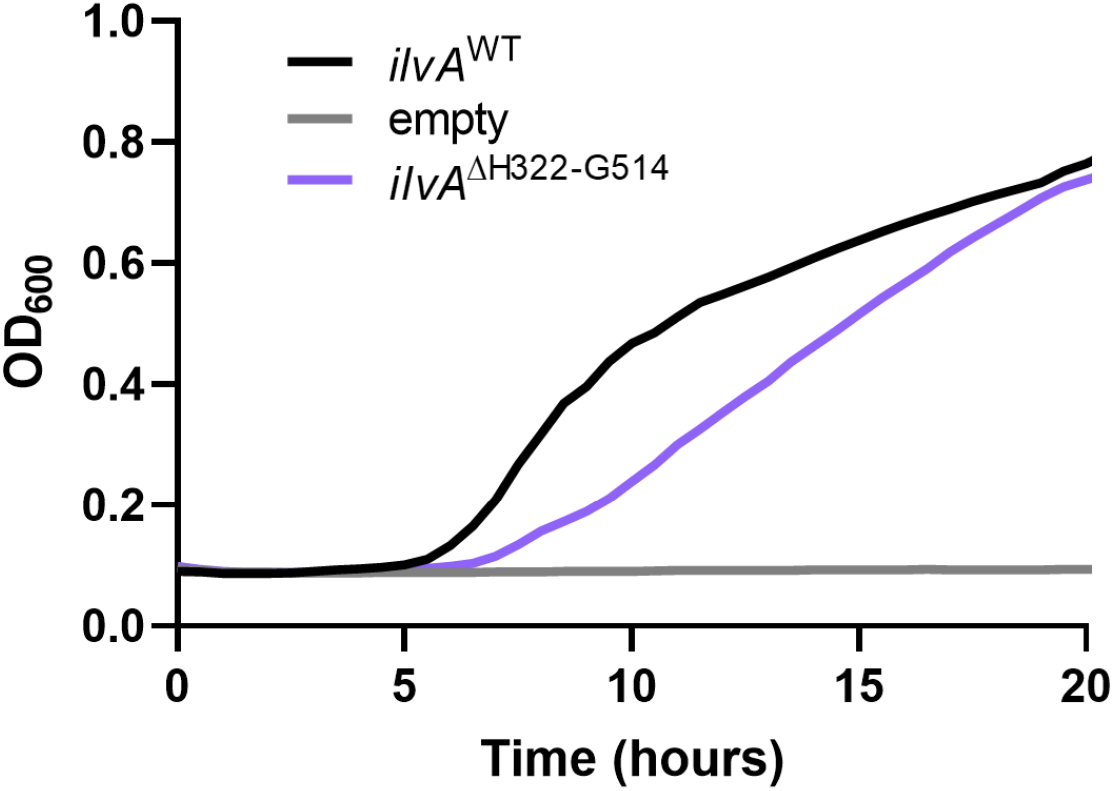
The C-terminus of *ilvA* is not required for isoleucine production in *P. protegens* Pf-5. A Δ*ilvAΔtdcB* P_*cymR*_-*relB P. protegens* strain harboring a vector containing the listed alleles under a rhamnose inducible promoter were grown in minimal medium without isoleucine and with rhamnose to probe the function of *ilvA*. The *ilvA*^ΔH322-G514^ allele contains a C-terminal truncation of *ilvA* immediately upstream of the internal RBS for *relE*. Growth is reported as OD_600_ over time. Data are shown as the mean of at least 3 independent replicates.

**Fig S8.**
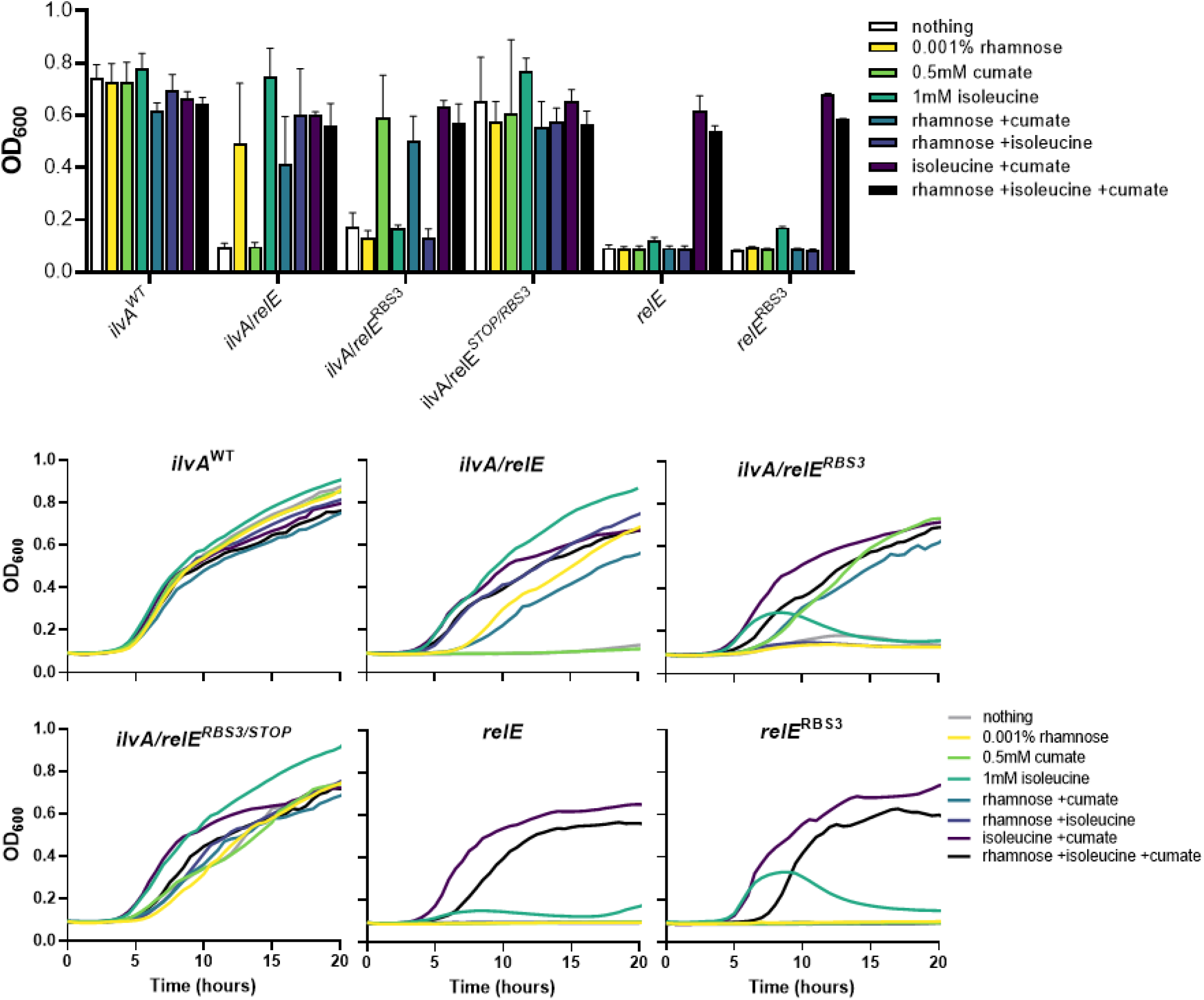
The *ilvA*/*relE*^*RBS3*^ and *relE*^*RBS3*^ alleles impart a growth defect in the absence of the antitoxin. Final density of isoleucine auxotroph (Δ*ilvAΔtdcB)* strains with chromosomally integrated antitoxin (P_*cymR*_-*relB*) and harboring plasmids carrying the different alleles listed under the P_*rhaBAD*_ promoter. **(A)** Strains harboring *ilvA*/*relE*^*STOP*^ alleles containing different strength RBSs were grown in minimal medium with addition of inducers as listed. Growth is reported as OD_600_ after 15 h. **(B)** Growth curves from data in panel **A** in which the OD_600_ was measured over time for each strain. Data in bar graphs are shown as the mean ± SD and data in growth curves are shown as the mean. All data are representative of 3 independent replicates.

**Fig S9.**
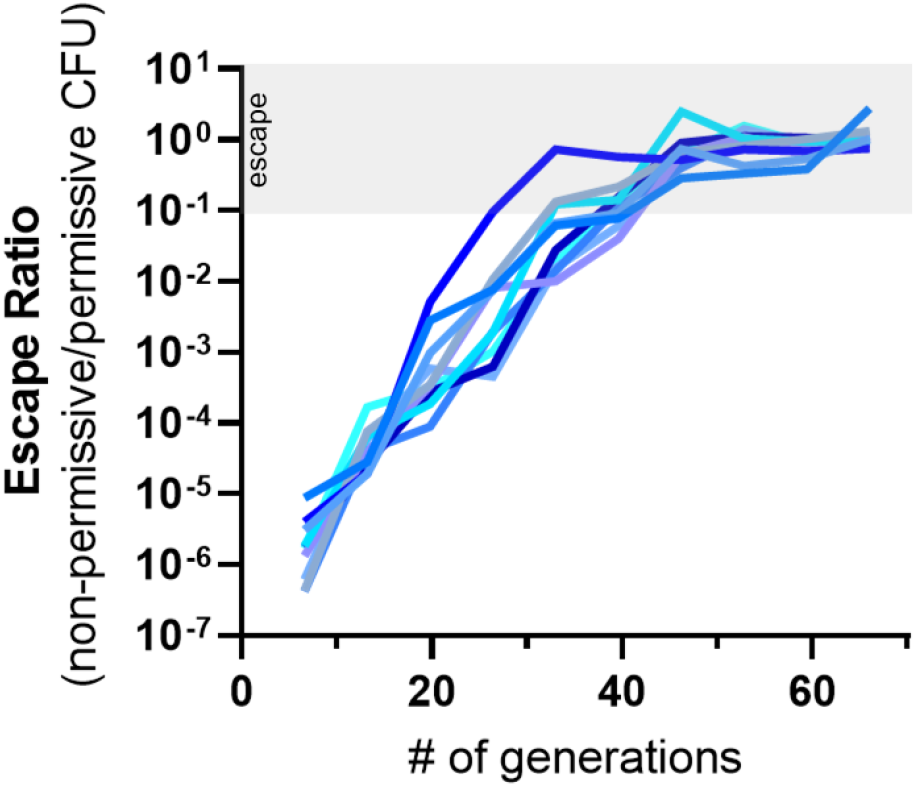
Evolutionary stability of non-entangled *relE*^RBS3^. Independent lineages (Δ*ilvA*Δ*tdcB* P_*cymR*_-*relB P. protegens*) harboring *relE*^*RBS3*^ were grown in minimal medium with rhamnose (to induce *relE*^RBS3^) and cumate (to induce antitoxin) in the presence of isoleucine. Each day (∼6.6 generations) the cultures were diluted 1:1,000 in fresh medium and plated for CFU on toxin permissive and non-permissive conditions and a survival frequency was calculated. The grey bar indicates toxin escape ratio ≥ 10^−1^ (10%). Each line represents data from one of ten independent replicates picked from single colonies. The RBS3 modification was engineered to drive the expression of P_*rham*_-*relE* to a similar level as *ilvA*/*relE*^*RBS3*^ (**Fig. S8**).

**Fig S10.**
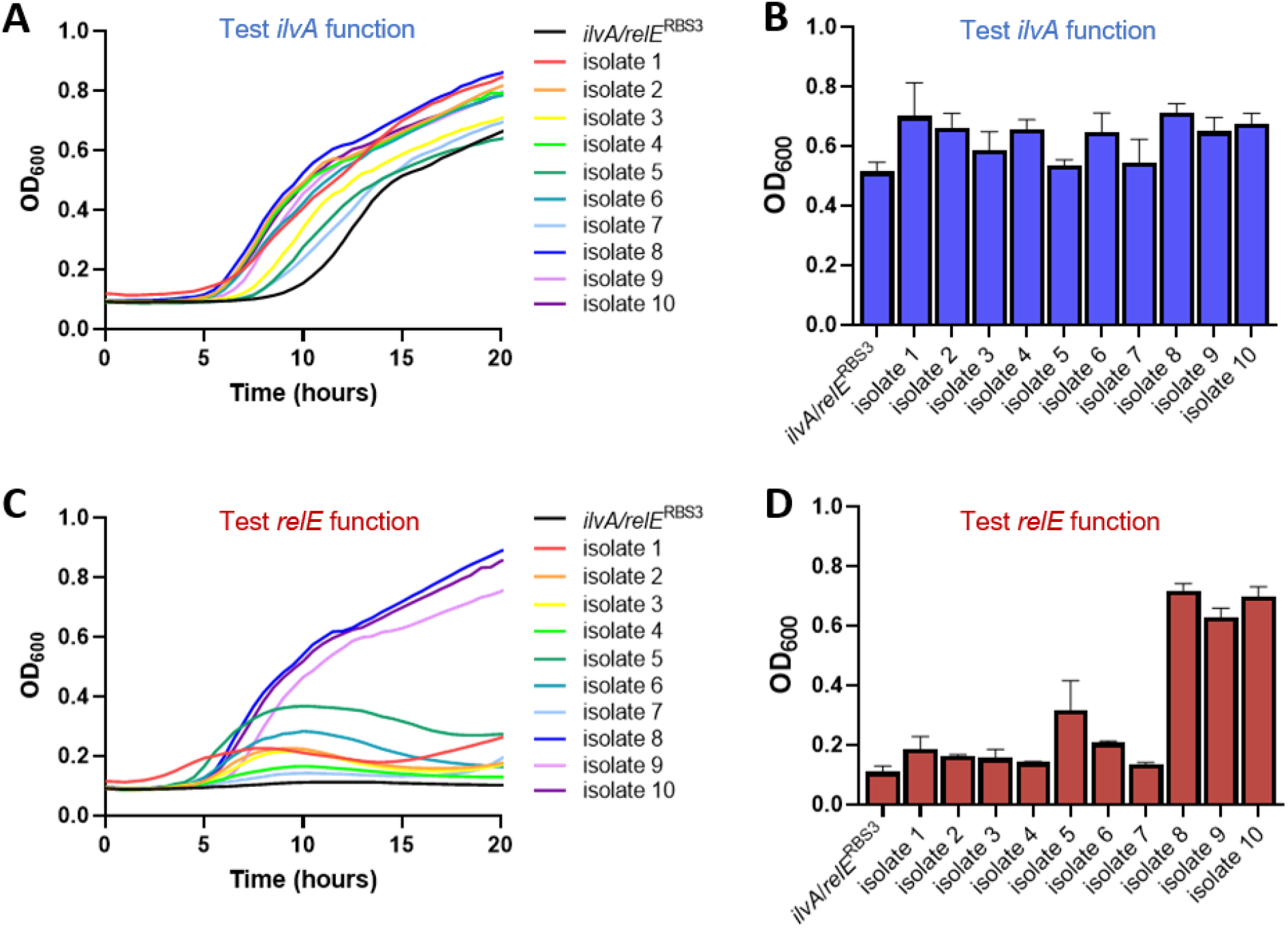
Growth of isolated colonies from each lineage of the long-term evolutionary stability assay. Single colonies were isolated from each lineage grown without isoleucine on permissive medium during the final passage of the long-term evolutionary stability assay. Each original colony was grown in minimal medium and the OD_600_ was measured over time (panels **A** and **C**) and a bar graph representing OD_600_ after 15 h of growth was generated (panels **B** and **D**). **(A, B)** To assess isoleucine auxotrophy, minimal medium without isoleucine was supplemented with rhamnose and cumate. **(C, D)** To assess *relE* toxicity, minimal medium was supplemented with rhamnose and isoleucine and without cumate. Data in bar graphs are shown as the mean ± SD and data in growth curves are shown as the mean. All data are representative of 3 independent replicates.

**Fig S11.**
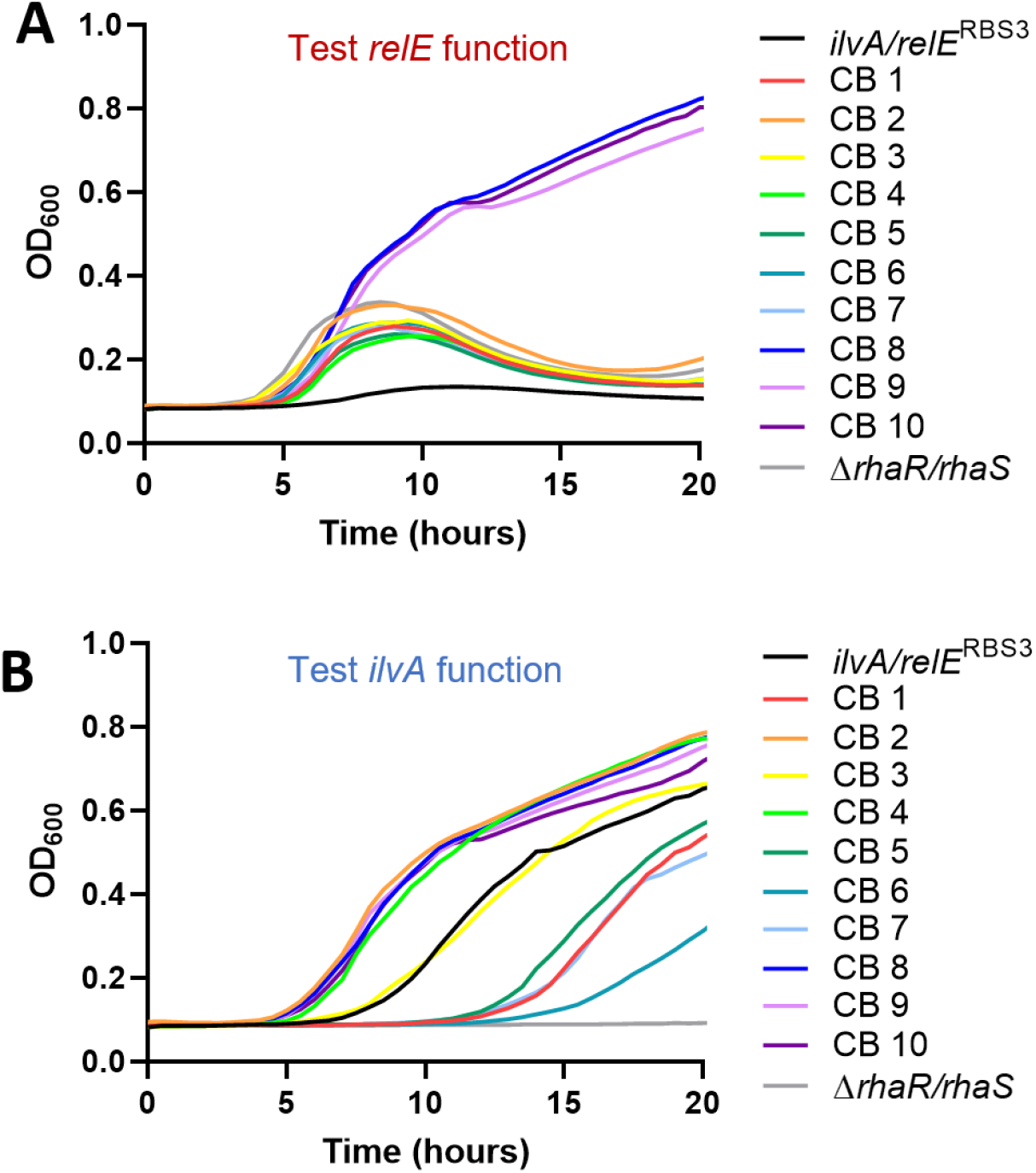
Growth of new clean background (CB) strains harboring isolated vectors from each lineage of the long-term evolutionary stability assay. Vectors were isolated from the colonies of each lineage grown without isoleucine and transformed into a clean genetic background (Δ*ilvAΔtdcB P*_*cymR*_*-relB)* and were grown in (**A**) minimal medium with isoleucine and rhamnose and without cumate to test *relE* activity or (**B**) in minimal medium without isoleucine with addition of rhamnose and cumate to probe *ilvA* function. Growth curves represent data from figure **4B** and **4C**. Data are shown as the mean of 3 independent replicates.

**Fig S12.**
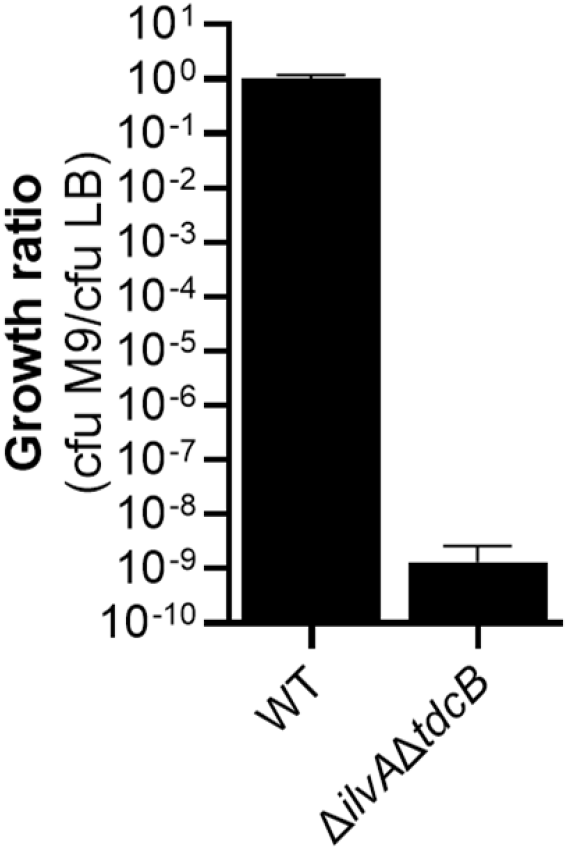
Growth ratio of cells that can survive on minimal medium. The indicated strains were grown overnight in LB medium, washed twice in minimal medium and then serially diluted onto either LB-agar or M9-agar plates without isoleucine. After 48 h of growth, growth ratio was calculated by taking the CFU/mL on M9 plates divided by the CFU/mL on LB plates. No colonies were observed for one replicate for the Δ*ilvA*Δ*tdcB* strain and this was reported as a limit of detection (one colony) at the lowest dilution plated. Data are shown as the mean ± SD of 4 independent replicates.

**Table S1.**
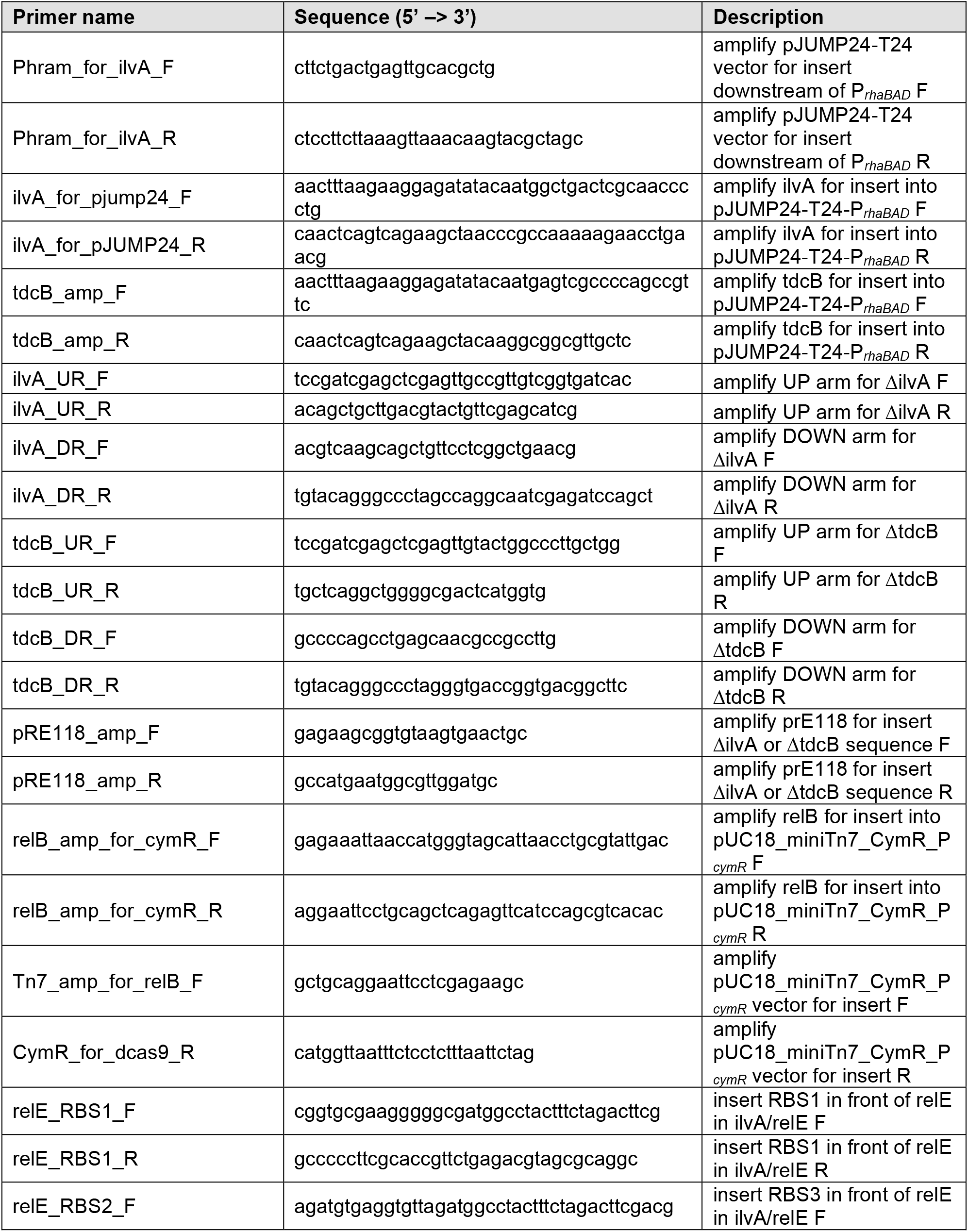
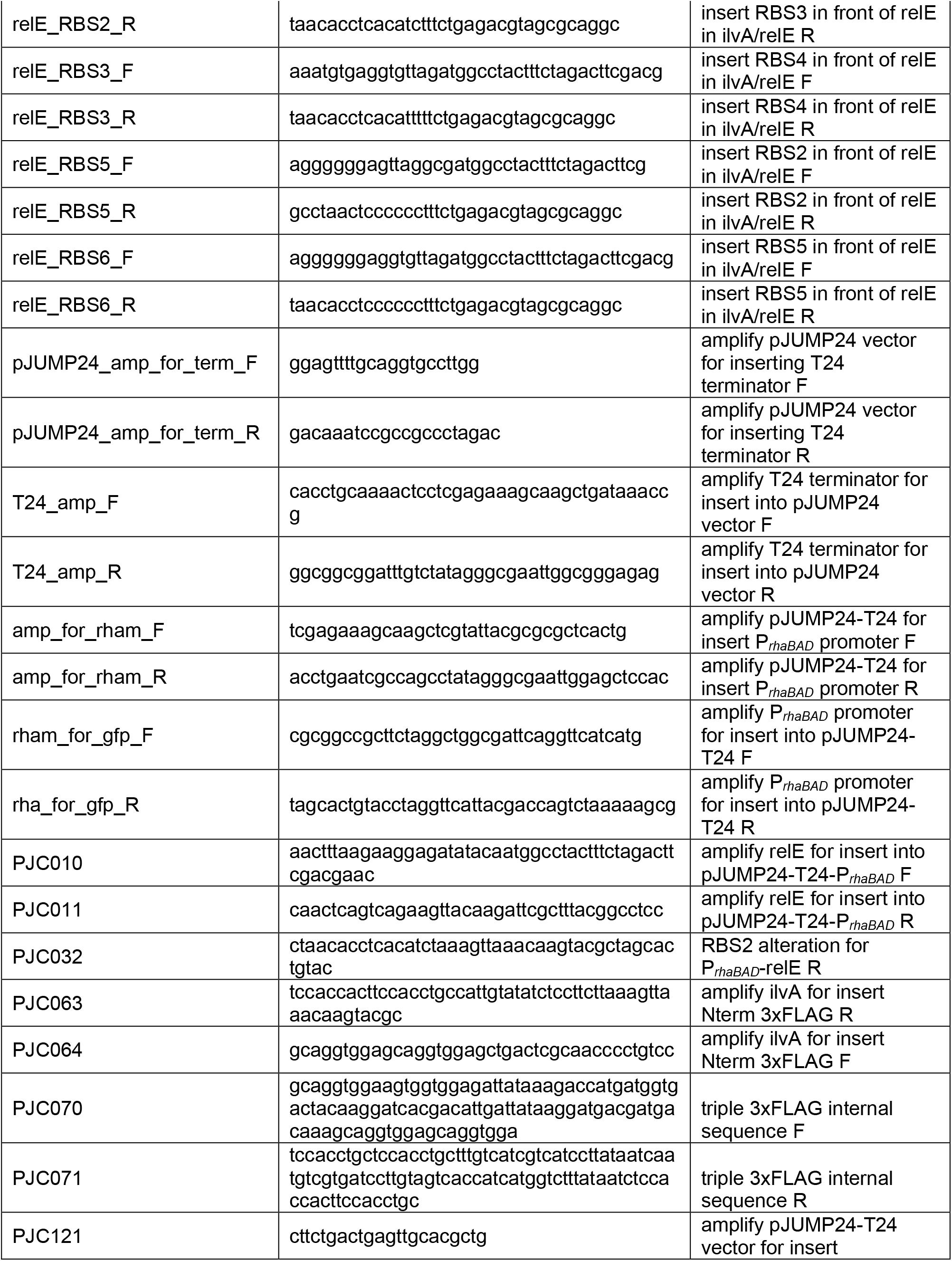
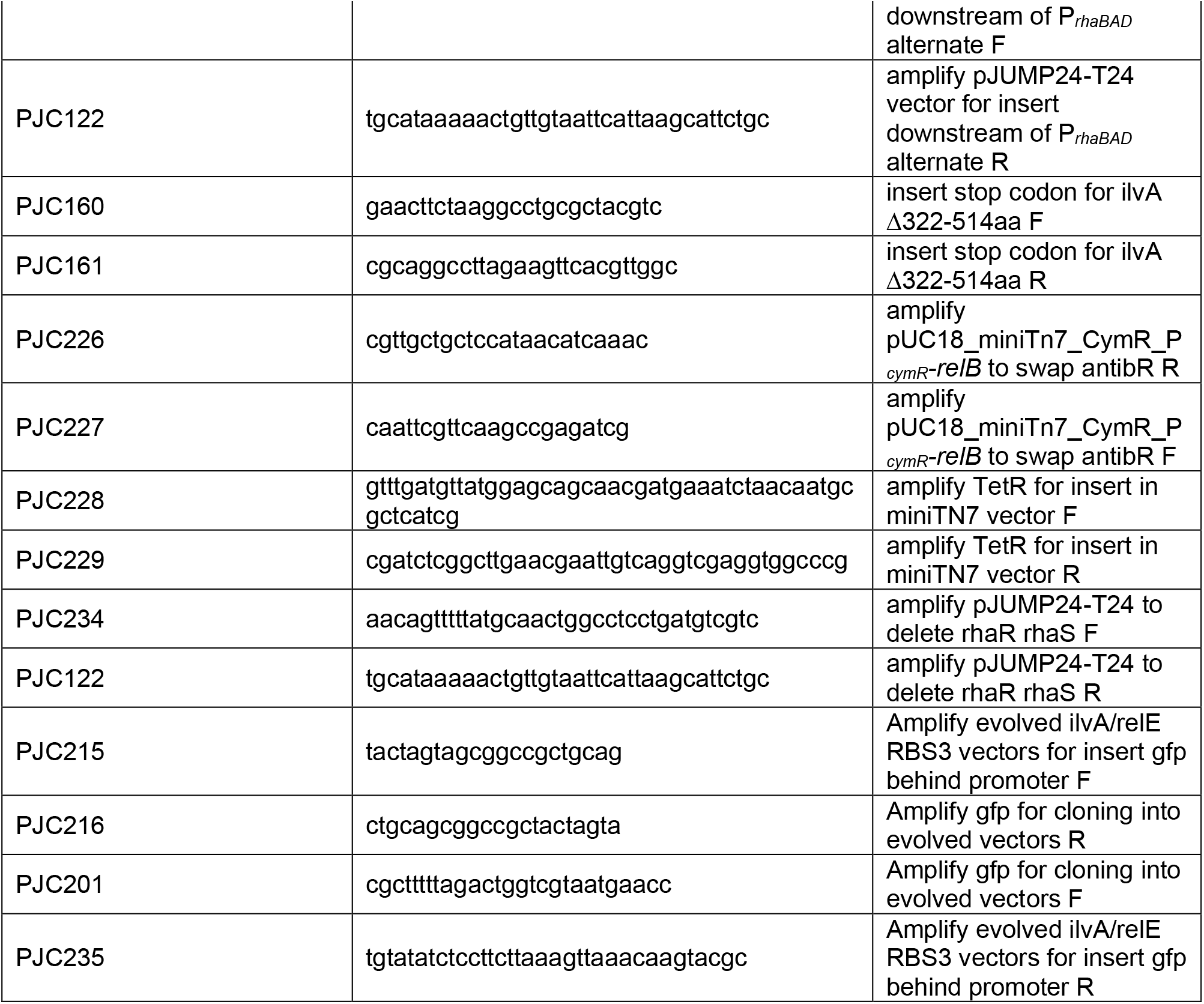
Primers used in this study.

**Table S2.**
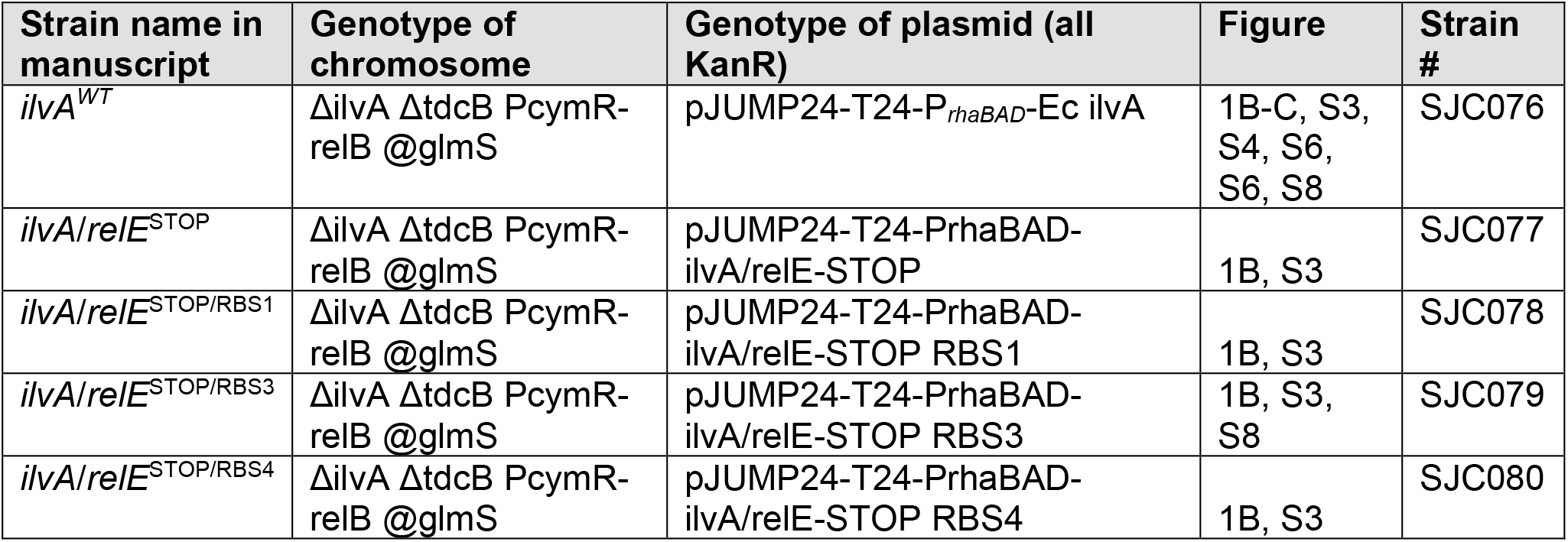
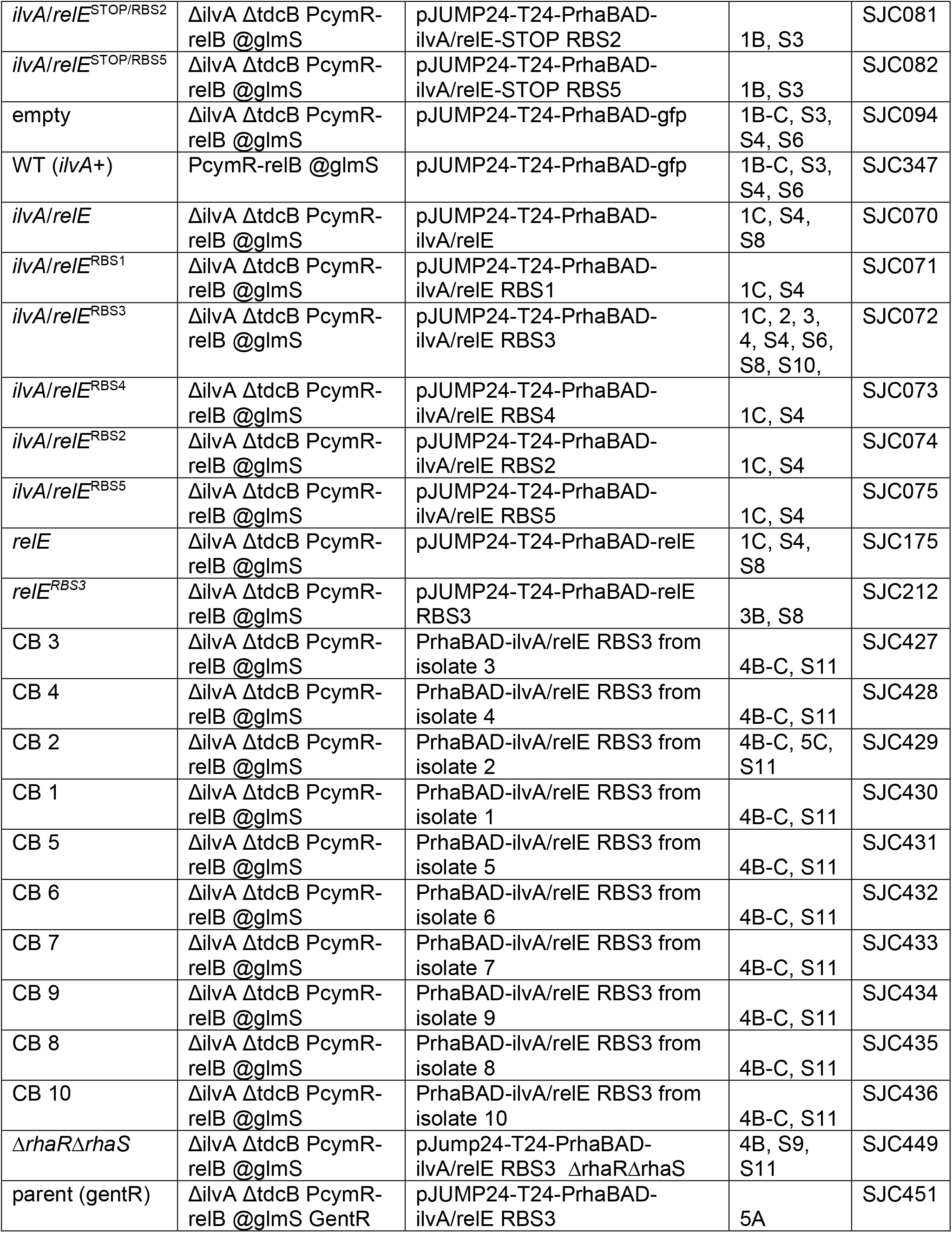
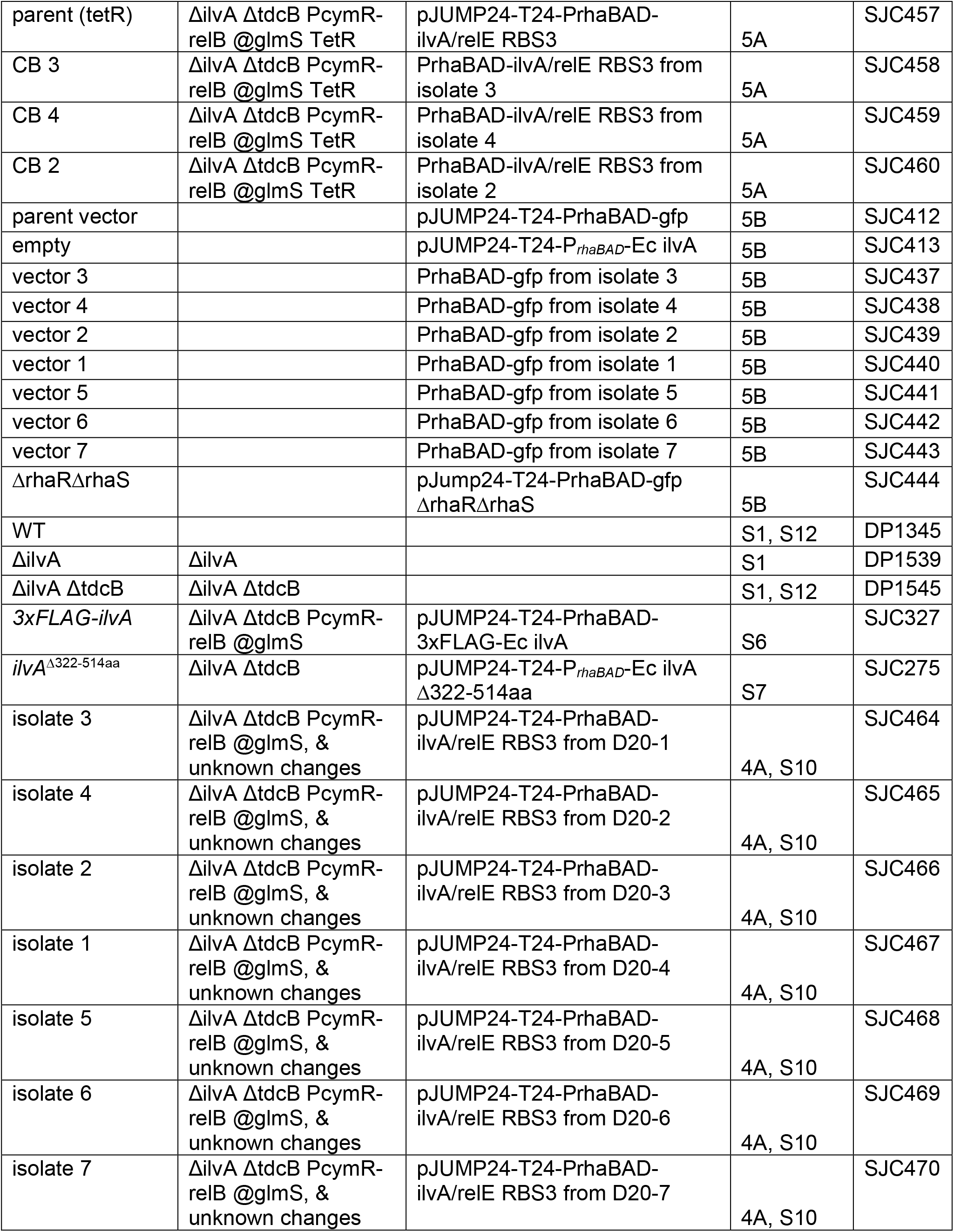
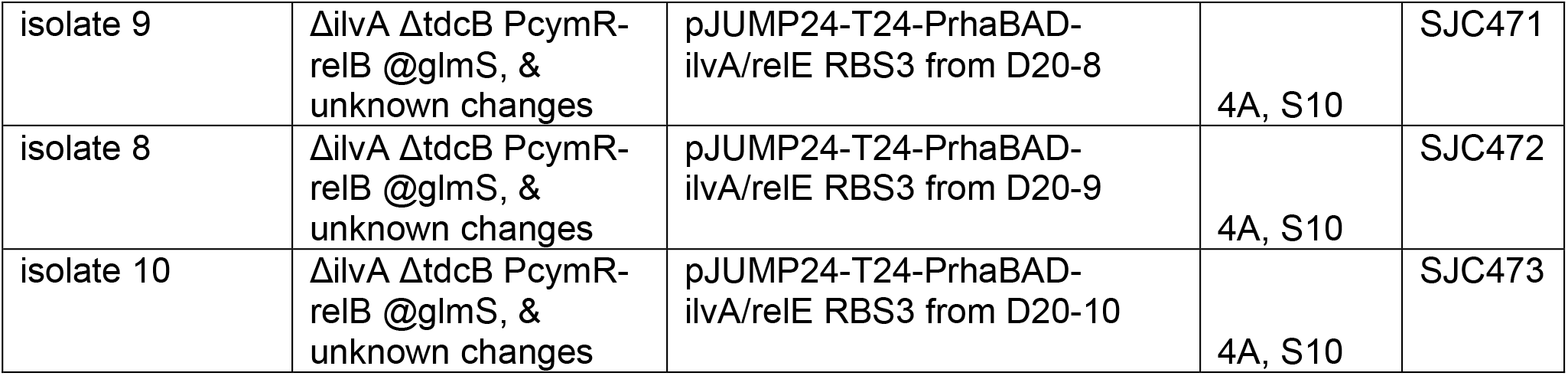
*Pseudomonas protegens* Pf-5 strains generated and used in this study.

| Strain name in manuscript | Genotype of chromosome | Genotype of plasmid (all KanR) | Figure | Strain # |
| --- | --- | --- | --- | --- |
| *ilvA<sup>WT</sup>* | Δ*ilvA* Δ*tdcB* *PcymR-reIB @glmS* | pJUMP24-T24-*P<sub>rhaBAD</sub>*-Ec *ilvA* | 1B-C, S3, S4, S6, S6, S8 | SJC076 |
| *ilvA/reIE<sup>STOP</sup>* | Δ*ilvA* Δ*tdcB* *PcymR-reIB @glmS* | pJUMP24-T24-*PrhaBAD-ilvA/reIE-STOP* | 1B, S3 | SJC077 |
| *ilvA/reIE<sup>STOP/RBS1</sup>* | Δ*ilvA* Δ*tdcB* *PcymR-reIB @glmS* | pJUMP24-T24-*PrhaBAD-ilvA/reIE-STOP RBS1* | 1B, S3 | SJC078 |
| *ilvA/reIE<sup>STOP/RBS3</sup>* | Δ*ilvA* Δ*tdcB* *PcymR-reIB @glmS* | pJUMP24-T24-*PrhaBAD-ilvA/reIE-STOP RBS3* | 1B, S3, S8 | SJC079 |
| *ilvA/reIE<sup>STOP/RBS4</sup>* | Δ*ilvA* Δ*tdcB* *PcymR-reIB @glmS* | pJUMP24-T24-*PrhaBAD-ilvA/reIE-STOP RBS4* | 1B, S3 | SJC080 |
| *ilvA/relE*<sup>STOP/RBS2</sup> | $\Delta ilvA \Delta tdcB$ PcymR-relB @glmS | pJUMP24-T24-PrhaBAD-ilvA/relE-STOP RBS2 | 1B, S3 | SJC081 |
| *ilvA/relE*<sup>STOP/RBS5</sup> | $\Delta ilvA \Delta tdcB$ PcymR-relB @glmS | pJUMP24-T24-PrhaBAD-ilvA/relE-STOP RBS5 | 1B, S3 | SJC082 |
| empty | $\Delta ilvA \Delta tdcB$ PcymR-relB @glmS | pJUMP24-T24-PrhaBAD-gfp | 1B-C, S3, S4, S6 | SJC094 |
| WT (*ilvA*+) ) | PcymR-relB @glmS | pJUMP24-T24-PrhaBAD-gfp | 1B-C, S3, S4, S6 | SJC347 |
| *ilvA/relE* | $\Delta ilvA \Delta tdcB$ PcymR-relB @glmS | pJUMP24-T24-PrhaBAD-ilvA/relE | 1C, S4, S8 | SJC070 |
| *ilvA/relE*<sup>RBS1</sup> | $\Delta ilvA \Delta tdcB$ PcymR-relB @glmS | pJUMP24-T24-PrhaBAD-ilvA/relE RBS1 | 1C, S4 | SJC071 |
| *ilvA/relE*<sup>RBS3</sup> | $\Delta ilvA \Delta tdcB$ PcymR-relB @glmS | pJUMP24-T24-PrhaBAD-ilvA/relE RBS3 | 1C, 2, 3, 4, S4, S6, S8, S10, | SJC072 |
| *ilvA/relE*<sup>RBS4</sup> | $\Delta ilvA \Delta tdcB$ PcymR-relB @glmS | pJUMP24-T24-PrhaBAD-ilvA/relE RBS4 | 1C, S4 | SJC073 |
| *ilvA/relE*<sup>RBS2</sup> | $\Delta ilvA \Delta tdcB$ PcymR-relB @glmS | pJUMP24-T24-PrhaBAD-ilvA/relE RBS2 | 1C, S4 | SJC074 |
| *ilvA/relE*<sup>RBS5</sup> | $\Delta ilvA \Delta tdcB$ PcymR-relB @glmS | pJUMP24-T24-PrhaBAD-ilvA/relE RBS5 | 1C, S4 | SJC075 |
| *relE* | $\Delta ilvA \Delta tdcB$ PcymR-relB @glmS | pJUMP24-T24-PrhaBAD-relE | 1C, S4, S8 | SJC175 |
| *relE*<sup>RBS3</sup> | $\Delta ilvA \Delta tdcB$ PcymR-relB @glmS | pJUMP24-T24-PrhaBAD-relE RBS3 | 3B, S8 | SJC212 |
| CB 3 | $\Delta ilvA \Delta tdcB$ PcymR-relB @glmS | PrhaBAD-ilvA/relE RBS3 from isolate 3 | 4B-C, S11 | SJC427 |
| CB 4 | $\Delta ilvA \Delta tdcB$ PcymR-relB @glmS | PrhaBAD-ilvA/relE RBS3 from isolate 4 | 4B-C, S11 | SJC428 |
| CB 2 | $\Delta ilvA \Delta tdcB$ PcymR-relB @glmS | PrhaBAD-ilvA/relE RBS3 from isolate 2 | 4B-C, 5C, S11 | SJC429 |
| CB 1 | $\Delta ilvA \Delta tdcB$ PcymR-relB @glmS | PrhaBAD-ilvA/relE RBS3 from isolate 1 | 4B-C, S11 | SJC430 |
| CB 5 | $\Delta ilvA \Delta tdcB$ PcymR-relB @glmS | PrhaBAD-ilvA/relE RBS3 from isolate 5 | 4B-C, S11 | SJC431 |
| CB 6 | $\Delta ilvA \Delta tdcB$ PcymR-relB @glmS | PrhaBAD-ilvA/relE RBS3 from isolate 6 | 4B-C, S11 | SJC432 |
| CB 7 | $\Delta ilvA \Delta tdcB$ PcymR-relB @glmS | PrhaBAD-ilvA/relE RBS3 from isolate 7 | 4B-C, S11 | SJC433 |
| CB 9 | $\Delta ilvA \Delta tdcB$ PcymR-relB @glmS | PrhaBAD-ilvA/relE RBS3 from isolate 9 | 4B-C, S11 | SJC434 |
| CB 8 | $\Delta ilvA \Delta tdcB$ PcymR-relB @glmS | PrhaBAD-ilvA/relE RBS3 from isolate 8 | 4B-C, S11 | SJC435 |
| CB 10 | $\Delta ilvA \Delta tdcB$ PcymR-relB @glmS | PrhaBAD-ilvA/relE RBS3 from isolate 10 | 4B-C, S11 | SJC436 |
| $\Delta rhaR \Delta rhaS$ | $\Delta ilvA \Delta tdcB$ PcymR-relB @glmS | pJump24-T24-PrhaBAD-ilvA/relE RBS3 $\Delta rhaR \Delta rhaS$ | 4B, S9, S11 | SJC449 |
| parent (gentR) | $\Delta ilvA \Delta tdcB$ PcymR-relB @glmS GentR | pJUMP24-T24-PrhaBAD-ilvA/relE RBS3 | 5A | SJC451 |
| parent (tetR) | $\Delta ilvA \Delta tdcB$ PcymR-relB @glmS TetR | pJUMP24-T24-PrhaBAD-ilvA/relE RBS3 | 5A | SJC457 |
| CB 3 | $\Delta ilvA \Delta tdcB$ PcymR-relB @glmS TetR | PrhaBAD-ilvA/relE RBS3 from isolate 3 | 5A | SJC458 |
| CB 4 | $\Delta ilvA \Delta tdcB$ PcymR-relB @glmS TetR | PrhaBAD-ilvA/relE RBS3 from isolate 4 | 5A | SJC459 |
| CB 2 | $\Delta ilvA \Delta tdcB$ PcymR-relB @glmS TetR | PrhaBAD-ilvA/relE RBS3 from isolate 2 | 5A | SJC460 |
| parent vector |  | pJUMP24-T24-PrhaBAD-gfp | 5B | SJC412 |
| empty |  | pJUMP24-T24-P<sub>rhaBAD</sub>-Ec ilvA | 5B | SJC413 |
| vector 3 |  | PrhaBAD-gfp from isolate 3 | 5B | SJC437 |
| vector 4 |  | PrhaBAD-gfp from isolate 4 | 5B | SJC438 |
| vector 2 |  | PrhaBAD-gfp from isolate 2 | 5B | SJC439 |
| vector 1 |  | PrhaBAD-gfp from isolate 1 | 5B | SJC440 |
| vector 5 |  | PrhaBAD-gfp from isolate 5 | 5B | SJC441 |
| vector 6 |  | PrhaBAD-gfp from isolate 6 | 5B | SJC442 |
| vector 7 |  | PrhaBAD-gfp from isolate 7 | 5B | SJC443 |
| $\Delta rhaR \Delta rhaS$ | | pJump24-T24-PrhaBAD-gfp $\Delta rhaR \Delta rhaS$ | 5B | SJC444 |
| WT |  |  | S1, S12 | DP1345 |
| $\Delta ilvA$ | $\Delta ilvA$ | | S1 | DP1539 |
| $\Delta ilvA \Delta tdcB$ | $\Delta ilvA \Delta tdcB$ | | S1, S12 | DP1545 |
| 3xFLAG-ilvA | $\Delta ilvA \Delta tdcB$ PcymR-relB @glmS | pJUMP24-T24-PrhaBAD-3xFLAG-Ec ilvA | S6 | SJC327 |
| *ilvA*<sup>$\Delta 322-514aa$</sup> | $\Delta ilvA \Delta tdcB$ | pJUMP24-T24-P<sub>rhaBAD</sub>-Ec ilvA $\Delta 322-514aa$ | S7 | SJC275 |
| isolate 3 | $\Delta ilvA \Delta tdcB$ PcymR-relB @glmS, & unknown changes | pJUMP24-T24-PrhaBAD-ilvA/relE RBS3 from D20-1 | 4A, S10 | SJC464 |
| isolate 4 | $\Delta ilvA \Delta tdcB$ PcymR-relB @glmS, & unknown changes | pJUMP24-T24-PrhaBAD-ilvA/relE RBS3 from D20-2 | 4A, S10 | SJC465 |
| isolate 2 | $\Delta ilvA \Delta tdcB$ PcymR-relB @glmS, & unknown changes | pJUMP24-T24-PrhaBAD-ilvA/relE RBS3 from D20-3 | 4A, S10 | SJC466 |
| isolate 1 | $\Delta ilvA \Delta tdcB$ PcymR-relB @glmS, & unknown changes | pJUMP24-T24-PrhaBAD-ilvA/relE RBS3 from D20-4 | 4A, S10 | SJC467 |
| isolate 5 | $\Delta ilvA \Delta tdcB$ PcymR-relB @glmS, & unknown changes | pJUMP24-T24-PrhaBAD-ilvA/relE RBS3 from D20-5 | 4A, S10 | SJC468 |
| isolate 6 | $\Delta ilvA \Delta tdcB$ PcymR-relB @glmS, & unknown changes | pJUMP24-T24-PrhaBAD-ilvA/relE RBS3 from D20-6 | 4A, S10 | SJC469 |
| isolate 7 | $\Delta ilvA \Delta tdcB$ PcymR-relB @glmS, & unknown changes | pJUMP24-T24-PrhaBAD-ilvA/relE RBS3 from D20-7 | 4A, S10 | SJC470 |
| isolate 9 | ΔilvA ΔtdcB PcymR-relB @glmS, & unknown changes | pJUMP24-T24-PrhaBAD-ilvA/relE RBS3 from D20-8 | 4A, S10 | SJC471 |
| isolate 8 | ΔilvA ΔtdcB PcymR-relB @glmS, & unknown changes | pJUMP24-T24-PrhaBAD-ilvA/relE RBS3 from D20-9 | 4A, S10 | SJC472 |
| isolate 10 | ΔilvA ΔtdcB PcymR-relB @glmS, & unknown changes | pJUMP24-T24-PrhaBAD-ilvA/relE RBS3 from D20-10 | 4A, S10 | SJC473 |

**Table S3.**
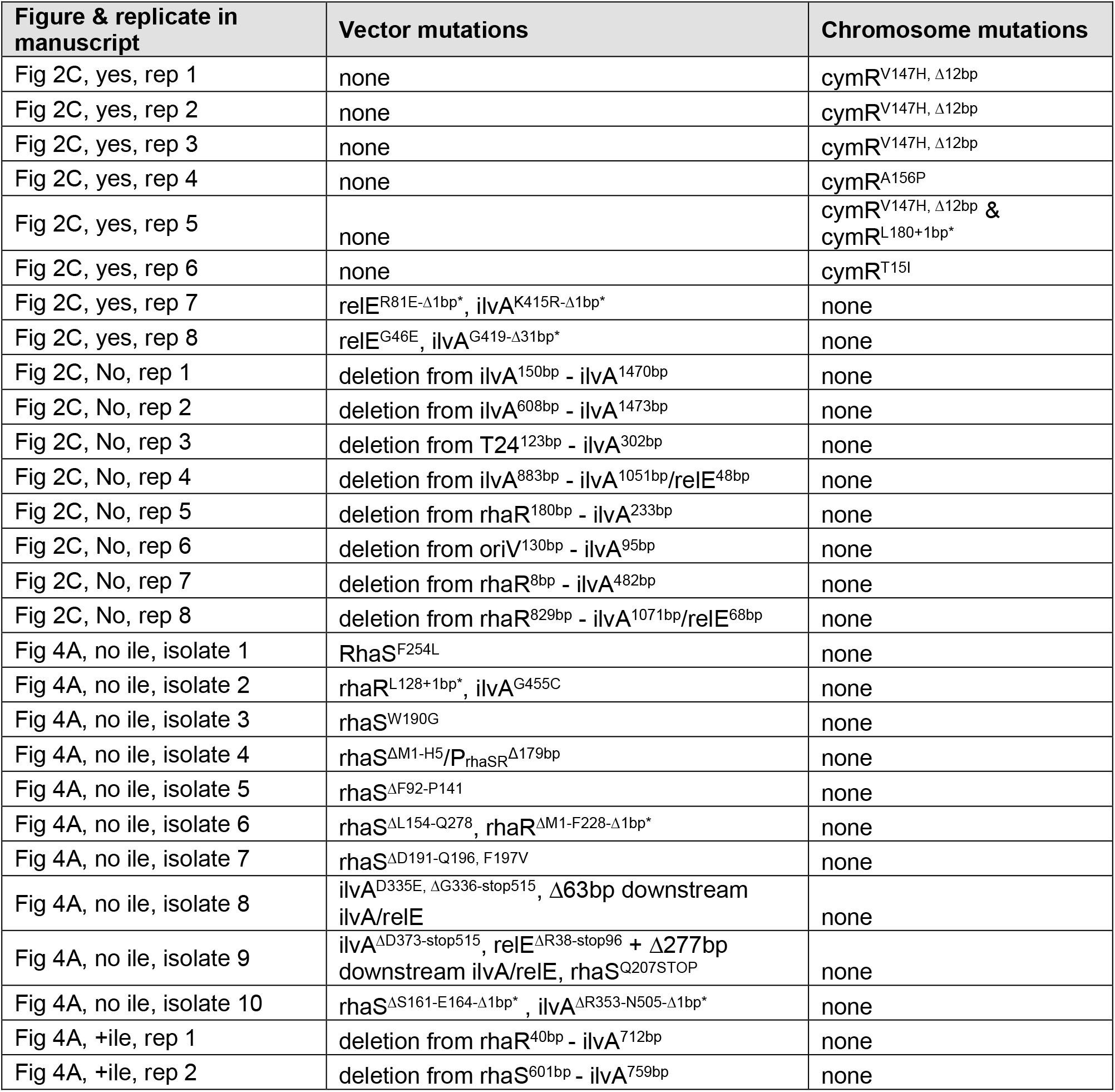
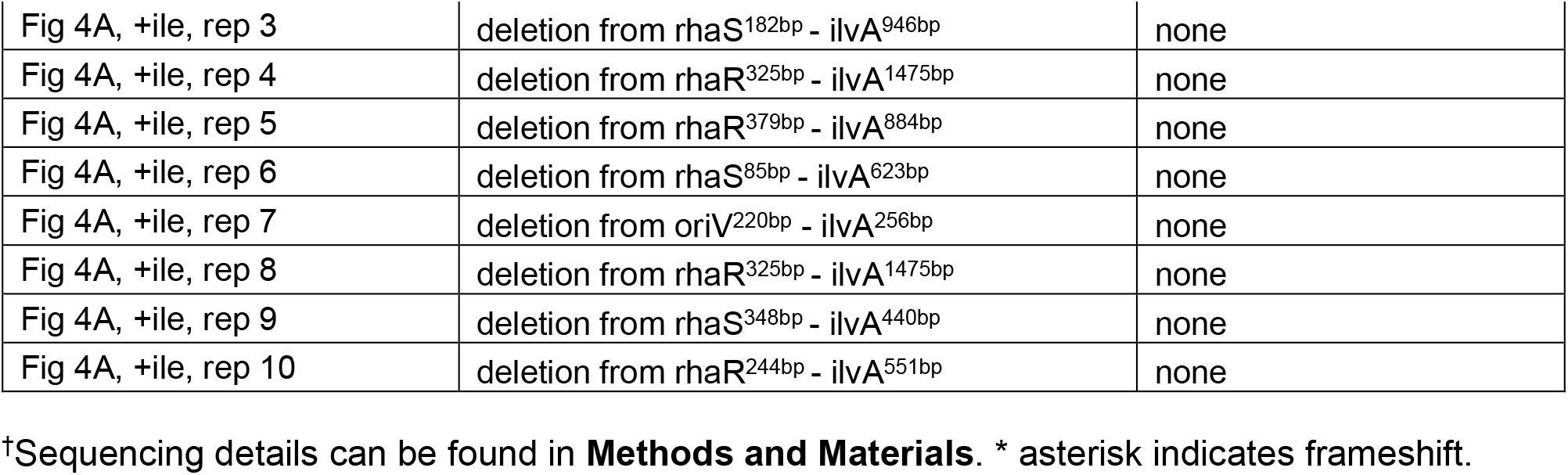
List of mutations in each sequenced strain of this study.

**Table S4.** Statistical Comparisons.

| Comparison | P value<sup>#</sup> |  | Information |
| --- | --- | --- | --- |
| Figure 2B - Paired Two-tailed Student's T-test |  |  |  |
| ilvA/reIE<sup>RBS3</sup>: + vs - isoleucine | *** | <0.0001 | Alpha = 0.05 |
| Figure 3B - One way ANOVA with Tukey's multiple comparison test |  |  |  |
| ilvA/reIE<sup>RBS3</sup> (- ile) vs. ilvA/reIE<sup>RBS3</sup> (+ ile) | **** | <0.0001 | # of families = 1 |
| ilvA/reIE<sup>RBS3</sup> (- ile) vs. reIE<sup>RBS3</sup> (+ ile) | **** | <0.0001 | # of comparisons per family = 3 |
| ilvA/reIE<sup>RBS3</sup> (+ ile) vs. reIE<sup>RBS3</sup> (+ ile) | ns | 0.1470 | Alpha = 0.05 |
| Figure 5A - One way ANOVA with Tukey's multiple comparison test |  |  |  |
| parent:parent vs. CB 3:parent | * | 0.0355 | # of families = 1 |
| parent:parent vs. CB 4:parent | * | 0.0239 | # of comparisons per family = 6 |
| parent:parent vs. CB 2:parent | ** | 0.0056 | Alpha = 0.05 |
| CB 3:parent vs. CB 4:parent | ns | 0.9956 |  |
| CB 3:parent vs. CB 2:parent | ns | 0.7216 |  |
| CB 4:parent vs. CB 2:parent | ns | 0.8403 |  |
| Figure 5B - One way ANOVA with Dunnett's multiple comparison test |  |  |  |
| parent vector vs. empty | **** | <0.0001 | # of families = 2 |
| parent vector vs. vector 3 | **** | <0.0001 | # of comparisons per family = 9 |
| parent vector vs. vector 4 | **** | <0.0001 | Alpha = 0.05 |
| parent vector vs. vector 2 | **** | <0.0001 |  |
| parent vector vs. vector 1 | **** | <0.0001 |  |
| parent vector vs. vector 5 | **** | <0.0001 |  |
| parent vector vs. vector 6 | **** | <0.0001 |  |
| parent vector vs. vector 7 | **** | <0.0001 |  |
| parent vector vs. ΔrhaRΔrhaS | **** | <0.0001 |  |
| empty vs. parent vector | **** | <0.0001 |  |
| empty vs. vector 3 | ** | 0.0020 |  |
| empty vs. vector 4 | ** | 0.0073 |  |
| empty vs. vector 2 | **** | <0.0001 |  |
| empty vs. vector 1 | ** | 0.0055 |  |
| empty vs. vector 5 | ** | 0.0034 |  |
| empty vs. vector 6 | * | 0.0113 |  |
| empty vs. vector 7 | ** | 0.0023 |  |
| empty vs. ΔrhaRΔrhaS | ** | 0.0045 | |

#Statistical differences were assessed using GraphPad Prism v.9.5.1 software. For escape ratio experiment in **Fig. 2B**, statistical analysis was performed on the log transformed data.

## REFERENCES

1. Sole, R.V., Montanez, R. and Duran-Nebreda, S. (2015) Synthetic circuit designs for earth terraformation. Biol Direct, 10, 37.

2. Puurunen, M.K., Vockley, J., Searle, S.L., Sacharow, S.J., Phillips, J.A., 3rd, Denney, W.S., Goodlett, B.D., Wagner, D.A., Blankstein, L., Castillo, M.J. et al. (2021) Safety and pharmacodynamics of an engineered E. coli Nissle for the treatment of phenylketonuria: a first-in-human phase 1/2a study. Nat Metab, 3, 1125–1132.

3. Chen, Z., Guo, L., Zhang, Y., Walzem, R.L., Pendergast, J.S., Printz, R.L., Morris, L.C., Matafonova, E., Stien, X., Kang, L. et al. (2014) Incorporation of therapeutically modified bacteria into gut microbiota inhibits obesity. J Clin Invest, 124, 3391–3406.

4. Steidler, L., Hans, W., Schotte, L., Neirynck, S., Obermeier, F., Falk, W., Fiers, W. and Remaut, E. (2000) Treatment of murine colitis by Lactococcus lactis secreting interleukin-10. Science, 289, 1352–1355.

5. Lagenaur, L.A., Sanders-Beer, B.E., Brichacek, B., Pal, R., Liu, X., Liu, Y., Yu, R., Venzon, D., Lee, P.P. and Hamer, D.H. (2011) Prevention of vaginal SHIV transmission in macaques by a live recombinant Lactobacillus. Mucosal Immunol, 4, 648–657.

6. Xiang, S., Fruehauf, J. and Li, C.J. (2006) Short hairpin RNA-expressing bacteria elicit RNA interference in mammals. Nat Biotechnol, 24, 697–702.

7. Ishikawa, M., Kojima, T. and Hori, K. (2021) Development of a Biocontained Toluene-Degrading Bacterium for Environmental Protection. Microbiol Spectr, 9, e0025921.

8. Zhang, H., Sun, X. and Dai, M. (2022) Improving crop drought resistance with plant growth regulators and rhizobacteria: Mechanisms, applications, and perspectives. Plant Commun, 3, 100228.

9. Shulse, C.N., Chovatia, M., Agosto, C., Wang, G., Hamilton, M., Deutsch, S., Yoshikuni, Y. and Blow, M.J. (2019) Engineered Root Bacteria Release Plant-Available Phosphate from Phytate. Appl Environ Microbiol, 85.

10. Suarez, R., Wong, A., Ramirez, M., Barraza, A., Orozco Mdel, C., Cevallos, M.A., Lara, M., Hernandez, G. and Iturriaga, G. (2008) Improvement of drought tolerance and grain yield in common bean by overexpressing trehalose-6-phosphate synthase in rhizobia. Mol Plant Microbe Interact, 21, 958–966.

11. You, L., Cox, R.S., 3rd, Weiss, R. and Arnold, F.H. (2004) Programmed population control by cell-cell communication and regulated killing. Nature, 428, 868–871.

12. Pitera, D.J., Paddon, C.J., Newman, J.D. and Keasling, J.D. (2007) Balancing a heterologous mevalonate pathway for improved isoprenoid production in Escherichia coli. Metab Eng, 9, 193–207.

13. Sleight, S.C. and Sauro, H.M. (2013) Visualization of evolutionary stability dynamics and competitive fitness of Escherichia coli engineered with randomized multigene circuits. ACS Synth Biol, 2, 519–528.

14. Ceroni, F., Algar, R., Stan, G.B. and Ellis, T. (2015) Quantifying cellular capacity identifies gene expression designs with reduced burden. Nat Methods, 12, 415–418.

15. Cardinale, S. and Arkin, A.P. (2012) Contextualizing context for synthetic biology--identifying causes of failure of synthetic biological systems. Biotechnol J, 7, 856–866.

16. Glick, B.R. (1995) Metabolic load and heterologous gene expression. Biotechnol Adv, 13, 247–261.

17. Lee, J.W., Chan, C.T.Y., Slomovic, S. and Collins, J.J. (2018) Next-generation biocontainment systems for engineered organisms. Nat Chem Biol, 14, 530–537.

18. Torres, L., Kruger, A., Csibra, E., Gianni, E. and Pinheiro, V.B. (2016) Synthetic biology approaches to biological containment: pre-emptively tackling potential risks. Essays Biochem, 60, 393–410.

19. Chan, C.T., Lee, J.W., Cameron, D.E., Bashor, C.J. and Collins, J.J. (2016) ‘Deadman’ and ‘Passcode’ microbial kill switches for bacterial containment. Nat Chem Biol, 12, 82–86.

20. Rottinghaus, A.G., Ferreiro, A., Fishbein, S.R.S., Dantas, G. and Moon, T.S. (2022) Genetically stable CRISPR-based kill switches for engineered microbes. Nat Commun, 13, 672.

21. Halvorsen, T.M., Ricci, D.P., Park, D.M., Jiao, Y. and Yung, M.C. (2022) Comparison of Kill Switch Toxins in Plant-Beneficial Pseudomonas fluorescens Reveals Drivers of Lethality, Stability, and Escape. ACS Synth Biol, 11, 3785–3796.

22. Balagadde, F.K., You, L., Hansen, C.L., Arnold, F.H. and Quake, S.R. (2005) Long-term monitoring of bacteria undergoing programmed population control in a microchemostat. Science, 309, 137–140.

23. Csorgo, B., Feher, T., Timar, E., Blattner, F.R. and Posfai, G. (2012) Low-mutation-rate, reduced-genome Escherichia coli: an improved host for faithful maintenance of engineered genetic constructs. Microb Cell Fact, 11, 11.

24. Jack, B.R., Leonard, S.P., Mishler, D.M., Renda, B.A., Leon, D., Suarez, G.A. and Barrick, J.E. (2015) Predicting the Genetic Stability of Engineered DNA Sequences with the EFM Calculator. ACS Synth Biol, 4, 939–943.

25. Rugbjerg, P., Sarup-Lytzen, K., Nagy, M. and Sommer, M.O.A. (2018) Synthetic addiction extends the productive life time of engineered Escherichia coli populations. Proc Natl Acad Sci U S A, 115, 2347–2352.

26. Posfai, G., Plunkett, G., 3rd, Feher, T., Frisch, D., Keil, G.M., Umenhoffer, K., Kolisnychenko, V., Stahl, B., Sharma, S.S., de Arruda, M. et al. (2006) Emergent properties of reduced-genome Escherichia coli. Science, 312, 1044–1046.

27. Chavez, A., Pruitt, B.W., Tuttle, M., Shapiro, R.S., Cecchi, R.J., Winston, J., Turczyk, B.M., Tung, M., Collins, J.J. and Church, G.M. (2018) Precise Cas9 targeting enables genomic mutation prevention. Proc Natl Acad Sci U S A, 115, 3669–3673.

28. Deatherage, D.E., Leon, D., Rodriguez, A.E., Omar, S.K. and Barrick, J.E. (2018) Directed evolution of Escherichia coli with lower-than-natural plasmid mutation rates. Nucleic Acids Res, 46, 9236–9250.

29. Cao, M., Tran, V.G. and Zhao, H. (2020) Unlocking nature’s biosynthetic potential by directed genome evolution. Curr Opin Biotechnol, 66, 95–104.

30. Yang, S., Sleight, S.C. and Sauro, H.M. (2013) Rationally designed bidirectional promoter improves the evolutionary stability of synthetic genetic circuits. Nucleic Acids Res, 41, e33.

31. Sleight, S.C., Bartley, B.A., Lieviant, J.A. and Sauro, H.M. (2010) Designing and engineering evolutionary robust genetic circuits. J Biol Eng, 4, 12.

32. Williams, R.L. and Murray, R.M. (2022) Integrase-mediated differentiation circuits improve evolutionary stability of burdensome and toxic functions in E. coli. Nat Commun, 13, 6822.

33. Liao, M.J., Din, M.O., Tsimring, L. and Hasty, J. (2019) Rock-paper-scissors: Engineered population dynamics increase genetic stability. Science, 365, 1045–1049.

34. Meyer, A.J., Segall-Shapiro, T.H., Glassey, E., Zhang, J. and Voigt, C.A. (2019) Escherichia coli “Marionette” strains with 12 highly optimized small-molecule sensors. Nat Chem Biol, 15, 196–204.

35. Blazejewski, T., Ho, H.I. and Wang, H.H. (2019) Synthetic sequence entanglement augments stability and containment of genetic information in cells. Science, 365, 595–598.

36. Wright, B.W., Molloy, M.P. and Jaschke, P.R. (2022) Overlapping genes in natural and engineered genomes. Nat Rev Genet, 23, 154–168.

37. Decrulle, A.L., Frenoy, A., Meiller-Legrand, T.A., Bernheim, A., Lotton, C., Gutierrez, A. and Lindner, A.B. (2021) Engineering gene overlaps to sustain genetic constructs in vivo. PLoS Comput Biol, 17, e1009475.

38. Miyata, T. and Yasunaga, T. (1978) Evolution of overlapping genes. Nature, 272, 532–535.

39. Simon-Loriere, E., Holmes, E.C. and Pagan, I. (2013) The effect of gene overlapping on the rate of RNA virus evolution. Mol Biol Evol, 30, 1916–1928.

40. Meydan, S., Marks, J., Klepacki, D., Sharma, V., Baranov, P.V., Firth, A.E., Margus, T., Kefi, A., Vazquez-Laslop, N. and Mankin, A.S. (2019) Retapamulin-Assisted Ribosome Profiling Reveals the Alternative Bacterial Proteome. Mol Cell, 74, 481–493 e486.

41. Meydan, S., Vazquez-Laslop, N. and Mankin, A.S. (2018) Genes within Genes in Bacterial Genomes. Microbiol Spectr, 6.

42. Wichmann, S., Scherer, S. and Ardern, Z. (2021) Biological factors in the synthetic construction of overlapping genes. BMC Genomics, 22, 888.

43. Lebre, S. and Gascuel, O. (2017) The combinatorics of overlapping genes. J Theor Biol, 415, 90–101.

44. Krakauer, D.C. (2000) Stability and evolution of overlapping genes. Evolution, 54, 731–739.

45. Safari, M., Jayaraman, B., Yang, S., Smith, C., Fernandes, J.D. and Frankel, A.D. (2022) Functional and structural segregation of overlapping helices in HIV-1. Elife, 11.

46. Ryu, M.H., Zhang, J., Toth, T., Khokhani, D., Geddes, B.A., Mus, F., Garcia-Costas, A., Peters, J.W., Poole, P.S., Ane, J.M. et al. (2020) Control of nitrogen fixation in bacteria that associate with cereals. Nat Microbiol, 5, 314–330.

47. Valenzuela-Ortega, M. and French, C. (2021) Joint universal modular plasmids (JUMP): a flexible vector platform for synthetic biology. Synth Biol (Oxf), 6, ysab003.

48. Choi, K.H. and Schweizer, H.P. (2006) mini-Tn7 insertion in bacteria with single attTn7 sites: example Pseudomonas aeruginosa. Nat Protoc, 1, 153–161.

49. Stephens, C., Reisenauer, A., Wright, R. and Shapiro, L. (1996) A cell cycle-regulated bacterial DNA methyltransferase is essential for viability. Proc. Natl. Acad. Sci. U.S.A., 93, 1210–1214.

50. Jeske, M. and Altenbuchner, J. (2010) The Escherichia coli rhamnose promoter rhaP(BAD) is in Pseudomonas putida KT2440 independent of Crp-cAMP activation. Appl Microbiol Biotechnol, 85, 1923–1933.

51. Klancher, C.A., Newman, J.D., Ball, A.S., van Kessel, J.C. and Dalia, A.B. (2020) Species-Specific Quorum Sensing Represses the Chitobiose Utilization Locus in Vibrio cholerae. Appl Environ Microbiol, 86.

52. Mavrodi, D.V., Loper, J.E., Paulsen, I.T. and Thomashow, L.S. (2009) Mobile genetic elements in the genome of the beneficial rhizobacterium Pseudomonas fluorescens Pf-5. BMC Microbiol, 9, 8.

53. Wright, O., Delmans, M., Stan, G.B. and Ellis, T. (2015) GeneGuard: A modular plasmid system designed for biosafety. ACS Synth Biol, 4, 307–316.

54. Umbarger, H.E. and Brown, B. (1956) Threonine deamination in Escherichia coli. I. D- and L-threonine deaminase activities of cell-free extracts. J Bacteriol, 71, 443–449.

55. Christensen, S.K. and Gerdes, K. (2003) RelE toxins from bacteria and Archaea cleave mRNAs on translating ribosomes, which are rescued by tmRNA. Mol Microbiol, 48, 1389–1400.

56. Pedersen, K., Zavialov, A.V., Pavlov, M.Y., Elf, J., Gerdes, K. and Ehrenberg, M. (2003) The bacterial toxin RelE displays codon-specific cleavage of mRNAs in the ribosomal A site. Cell, 112, 131–140.

57. Liang, Y.F., Long, Z.X., Zhang, Y.J., Luo, C.Y., Yan, L.T., Gao, W.Y. and Li, H. (2021) The chemical mechanisms of the enzymes in the branched-chain amino acids biosynthetic pathway and their applications. Biochimie, 184, 72–87.

58. Gallagher, D.T., Gilliland, G.L., Xiao, G., Zondlo, J., Fisher, K.E., Chinchilla, D. and Eisenstein, E. (1998) Structure and control of pyridoxal phosphate dependent allosteric threonine deaminase. Structure, 6, 465–475.

59. Hogan, A.M., Jeffers, K.R., Palacios, A. and Cardona, S.T. (2021) Improved Dynamic Range of a Rhamnose-Inducible Promoter for Gene Expression in Burkholderia spp. Appl Environ Microbiol, 87, e0064721.

60. Li, G.Y., Zhang, Y., Inouye, M. and Ikura, M. (2009) Inhibitory mechanism of Escherichia coli RelE-RelB toxin-antitoxin module involves a helix displacement near an mRNA interferase active site. J Biol Chem, 284, 14628–14636.

61. Eaton, R.W. (1996) p-Cumate catabolic pathway in Pseudomonas putida Fl: cloning and characterization of DNA carrying the cmt operon. J Bacteriol, 178, 1351–1362.

62. Eaton, R.W. (1997) p-Cymene catabolic pathway in Pseudomonas putida F1: cloning and characterization of DNA encoding conversion of p-cymene to p-cumate. J Bacteriol, 179, 3171–3180.

63. Halper, S.M., Hossain, A. and Salis, H.M. (2020) Synthesis Success Calculator: Predicting the Rapid Synthesis of DNA Fragments with Machine Learning. ACS Synth Biol, 9, 1563–1571.

64. Cetnar, D.P. and Salis, H.M. (2021) Systematic Quantification of Sequence and Structural Determinants Controlling mRNA stability in Bacterial Operons. ACS Synth Biol, 10, 318–332.

65. Bhende, P.M. and Egan, S.M. (1999) Amino acid-DNA contacts by RhaS: an AraC family transcription activator. J Bacteriol, 181, 5185–5192.

66. Hall, J.P.J., Wright, R.C.T., Harrison, E., Muddiman, K.J., Wood, A.J., Paterson, S. and Brockhurst, M.A. (2021) Plasmid fitness costs are caused by specific genetic conflicts enabling resolution by compensatory mutation. PLoS Biol, 19, e3001225.

67. Babu, M.M. and Aravind, L. (2006) Adaptive evolution by optimizing expression levels in different environments. Trends Microbiol, 14, 11–14.

68. Dekel, E. and Alon, U. (2005) Optimality and evolutionary tuning of the expression level of a protein. Nature, 436, 588–592.

69. Son, H.I., Weiss, A. and You, L. (2021) Design patterns for engineering genetic stability. Curr Opin Biomed Eng, 19.

70. Lenski, R.E. (2017) Experimental evolution and the dynamics of adaptation and genome evolution in microbial populations. ISME J, 11, 2181–2194.

71. Barrick, J.E. and Lenski, R.E. (2013) Genome dynamics during experimental evolution. Nat Rev Genet, 14, 827–839.

72. Stirling, F. and Silver, P.A. (2020) Controlling the Implementation of Transgenic Microbes: Are We Ready for What Synthetic Biology Has to Offer? Mol Cell, 78, 614–623.

73. Cotton, C.A., Bernhardsgrutter, I., He, H., Burgener, S., Schulz, L., Paczia, N., Dronsella, B., Erban, A., Toman, S., Dempfle, M. et al. (2020) Underground isoleucine biosynthesis pathways in E. coli. Elife, 9.

74. Liao, C., Wang, T., Maslov, S. and Xavier, J.B. (2020) Modeling microbial cross-feeding at intermediate scale portrays community dynamics and species coexistence. PLoS Comput Biol, 16, e1008135.

75. Mee, M.T., Collins, J.J., Church, G.M. and Wang, H.H. (2014) Syntrophic exchange in synthetic microbial communities. Proc Natl Acad Sci U S A, 111, E2149–2156.

76. Moe-Behrens, G.H., Davis, R. and Haynes, K.A. (2013) Preparing synthetic biology for the world. Front Microbiol, 4, 5.

77. Stirling, F., Bitzan, L., O’Keefe, S., Redfield, E., Oliver, J.W.K., Way, J. and Silver, P.A. (2017) Rational Design of Evolutionarily Stable Microbial Kill Switches. Mol Cell, 68, 686–697 e683.

## Supplemental Reference

1. Halper, S.M., Hossain, A. and Salis, H.M. (2020) Synthesis Success Calculator: Predicting the Rapid Synthesis of DNA Fragments with Machine Learning. ACS Synth Biol, 9, 1563-1571.

